# Attractor dynamics of a whole-cortex network model predicts emergence and structure of fMRI co-activation patterns in the mouse brain

**DOI:** 10.1101/2022.04.28.489908

**Authors:** Diego Fasoli, Ludovico Coletta, Daniel Gutierrez-Barragan, Alessandro Gozzi, Stefano Panzeri

## Abstract

Resting state fMRI activity in mammals exhibits rich dynamics on a fast, frame-by-frame timescale of seconds, including the robust emergence of recurring fMRI co-activation patterns (CAPs). To understand how such dynamics emerges from the underlying anatomical cortico-cortical connectivity, we developed a whole-cortex model of resting-state fMRI activity in the mouse. Our model implemented neural input-output nonlinearities and excitatory-inhibitory interactions within cortical regions, as well as directed anatomical connectivity between regions inferred from the Allen mouse brain atlas. We found that, even if the model parameters were fitted to explain static properties of fMRI activity on the timescale of minutes, the model generated rich frame-by-frame attractor dynamics, with multiple stationary and oscillatory attractors. Guided by these theoretical predictions, we found that empirical mouse fMRI time series exhibited analogous signatures of attractor dynamics, and that model attractors recapitulated the topographical organization of empirical fMRI CAPs. The model established key relationships between attractor dynamics, CAPs and features of the directed cortico-cortical intra- and inter-hemispheric anatomical connectivity. Specifically, we found that neglecting fiber directionality severely affected the number of model’s attractors and their ability to explain CAPs. Furthermore, modifying inter-hemispheric anatomical connectivity strength by decreasing or increasing it from the value of real mouse anatomical data, resulted in fewer attractors generated by cortico-cortical interactions and reduced non-homotopic features of the attractors generated by the model, which were important for better predicting empirical CAPs. These results offer novel theoretical insight into the dynamic organization of resting state fMRI in the mouse brain and suggest that the frame-wise BOLD activity captured by CAPs reflects an emerging property of cortical dynamics resulting from directed cortico-cortical interactions.

**Author summary:** Whole-brain activity at rest transitions on a timescale of seconds between stereotyped co-activation patterns (CAPs), each with a distinct spatial profile of co-activation across different brain regions. CAPs have been robustly reported across datasets and mammalian species, including humans and mice. However, the significance and origin of these patterns remain unknown. Here we studied the origin of CAPs using a computational model of the whole cortex based on real-world directed measurements of mouse cortical anatomical connectivity. We found that we could explain the formation and topography of CAPs in terms of attractors (that is, states the network tends to converge to) that reflect the information in the anatomical connections between cortical regions. Attractors and CAPs are significantly influenced by the directionality of connectivity (available in mice but not in humans) and the strength of inter-hemispheric coupling. Together, our findings provide a novel possible explanation of the mechanisms of CAP generation which relates to cortico-cortical connectivity, and suggest that CAPs are at least in part an emergent property of cortico-cortical interactions which can be generated also in absence of the action of arousing modulatory nuclei.

## 1 Introduction

Resting-state fMRI has been widely used to map functional organization of spontaneous large-scale activity in the human brain [1–3]. These measures have been paralleled and informed by computational models of large-scale interactions between regions of the brain. Such network models typically incorporate anatomical connections as inferred by diffusion tensor imaging, and are used to understand and predict the principles of the ensuing collective brain dynamics [4–7]. Computational models have initially been employed to describe steady-state measures of “static” functional connectivity (that is, averaged over many minutes of resting state activity [8–12]), and have more recently begun to be used to understand dynamical frame-by-frame changes in network configuration occurring on a timescale of individual fMRI time frames [13–19].

In recent years, fMRI studies in humans have been complemented by analogous neuroimaging investigations in animal species, including rodents. Mouse fMRI studies are important to understand the mechanisms and principles of large-scale brain activity, owing to the possibility of causally manipulating brain activity, or mimicking pathological states via genetic or environmental interventions [20–22]. The use of physiologically accessible species has also allowed a more precise measure of the underlying anatomical connectivity and its directionality [23–25], which cannot at present be estimated with analogous precision in humans owing to the limitation of Diffusion Tensor Imaging (DTI), a technique that lacks information on fiber directionality, and does not allow to reliably resolve long axonal tracts [26]. fMRI studies in mice have revealed important principles in the organization of neuronal activity, such as the formation of large-scale functional hubs and the relationship between functional and anatomical connectivity [23, 27–29]. Importantly, they have also revealed rich dynamics in mouse fMRI time series [30, 31], which partly recapitulates that seen in humans [32–35], and which is governed by a few dominant frame-by-frame co-activation patterns (CAPs) of fMRI activity across regions [35–37].

Because of the growing importance of these studies in the mouse, theoretical research has begun to extend network modelling to resting state fMRI timeseries acquired in this species [18, 38–40]. These models can help conceptualize the neurophysiological processes that shape resting state fMRI activity. For one, the mechanisms and principles underlying the emergence of CAPs remain elusive. Are CAPs generated by a subcortical executive, such as arousing modulatory nuclei, or are they a self-organizing phenomenon reflecting an emergent property of cortico-cortical interactions [41]? Addressing this issue with network models is of crucial importance given the emerging contribution of CAPs in shaping resting state fMRI dynamics in human and animals, and also the numerous and conflicting theories related to their origin [30, 31, 42–44].

To fill this knowledge gap, here we developed a whole-cortex neural network model that combines realistic directional anatomical connectivity of the mouse neocortex [23–25, 45] with non-linear firing rate dynamics in each region. The model allows us to simulate the emergent collective dynamics of the interconnected brain regions. We first fitted the model parameters to match the time-averaged pattern of relative activation across regions and the static functional connectivity matrix. We next used our ability to study in detail the dynamics of this network to make several novel predictions about the frame-by-frame dynamics of the whole-cortex resting state fMRI recordings and its relationship with the underlying anatomy of inter-regional connections. Specifically, we computed the attractor dynamics generated by the model, revealing a rich structure of oscillatory and stationary attractors. Model-assisted analysis of empirical data revealed that analogous signatures of attractor dynamics are present in empirical mouse resting state fMRI timeseries. We also found that features of real CAPs seen in mice can be explained by the model attractor dynamics and depend critically on the direction of anatomical wiring. The richness and complexity of attractor dynamics was maximal for values of the inter-hemispheric connection strength and of connectivity sparsity found in real anatomical mouse data.

## 2 Results

We designed a novel neural network model of the whole mouse cortex. The model was endowed with a directed matrix of the anatomical connectivity between cortical regions taken from mouse data, and it generated neural dynamics with plausible neural input-output non-linearities and excitatory-inhibitory interactions within each region. Our model generated fMRI signals from neural activity with a simple neurovascular coupling function, producing biologically plausible neural and resting-state fMRI dynamics for each region. Importantly, the models we designed was simple enough to allow for a thorough numerical and analytical study of its dynamics. All model parameters were selected to fit static resting-state fMRI activity and fMRI functional connectivity of real mouse data. The model was then used to make novel predictions about the role of specific elements of the anatomical connectivity in generating whole-cortex attractor dynamics. These model predictions were tested on real mouse resting state fMRI time series by comparing the predictions against frame-by-frame features of resting state fMRI functional connectivity such as CAPs.

### 2.1 Fitting a large-scale neural model of mouse cortex to time-averaged activity and functional connectivity of resting state fMRI data

We designed a large-scale neural network model (Fig. 1A) including 34 cortical regions of the mouse brain (17 per hemisphere). We chose to parcellate the cortex in 34 regions as this was the finest parcellation that still allowed us to study numerically and analytically in great detail attractor dynamics [46]. Each region was modelled to comprise one excitatory (E) and one inhibitory (I) mutually interacting neural populations. Each population was described as a threshold binary (active vs non-active) unit [47–49] whose states, representing the mean spiking activity of each population, were updated synchronously at discrete time steps. Each excitatory population within a region received inputs from both excitatory and inhibitory populations from the same region, as well as excitatory inputs from other cortical regions, whose strength was weighted by the anatomical connectivity matrix. Each inhibitory population received instead inhibitory and excitatory inputs from the same region, but not from other regions. Further, a noise term was added as input to both inhibitory and excitatory populations, to express the net effect of stochastic components of neural activity. As we focus on cortico-cortical interactions, we did not include inputs from subcortical regions.

**Figure 1:**
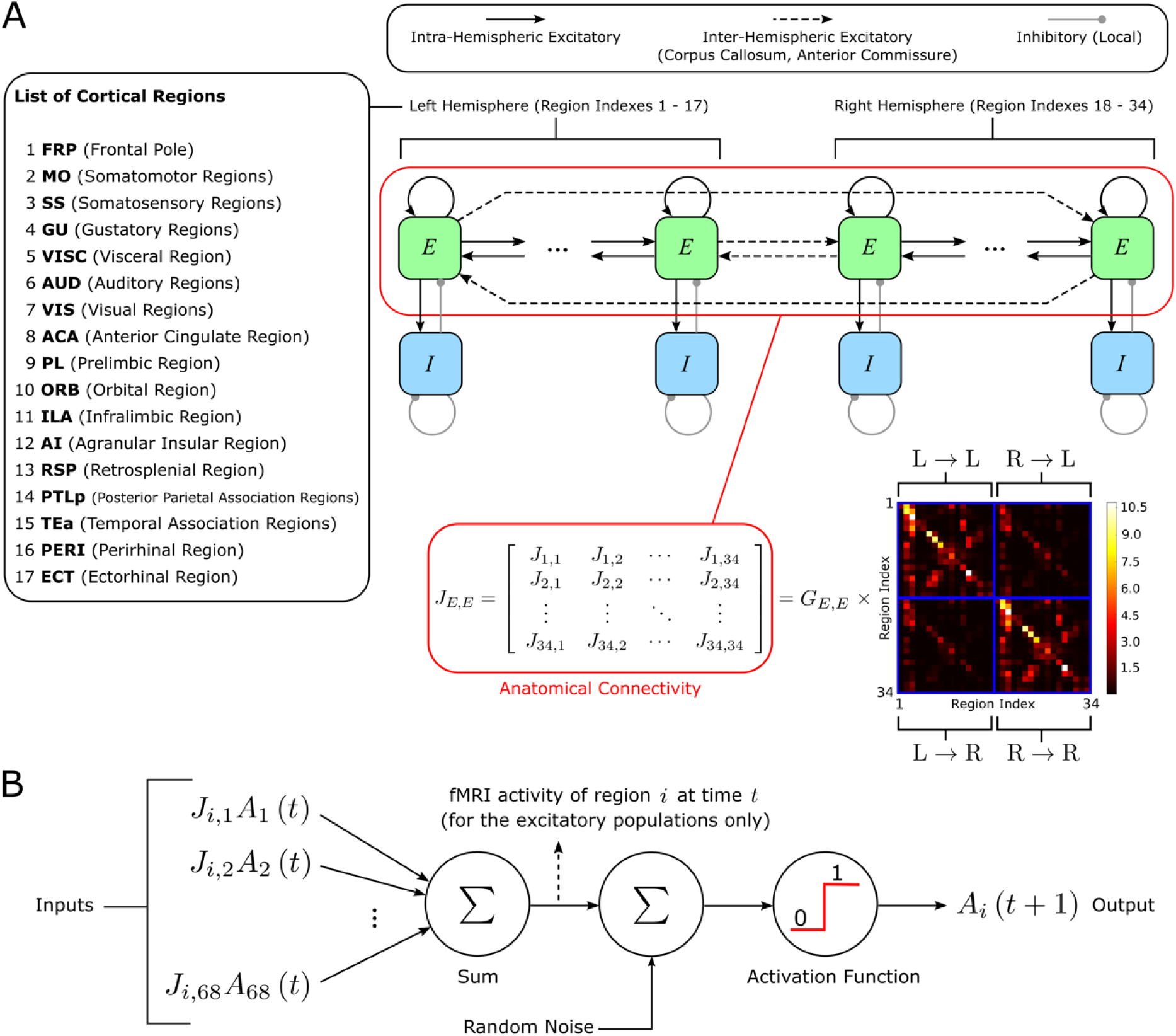
Model architecture. **A)** Our model incorporates 34 (17 per hemisphere) cortical regions (listed on the left), each with an excitatory (E) and an inhibitory (I) population. The excitatory populations across different regions are connected to each other through a directed anatomical connectivity matrix extracted from the mouse connectome (multiplied by a global scaling coefficient *G*_*E,E*_), and are also connected locally to the corresponding inhibitory population. **B)** Mathematical model that describes the temporal evolution of the spiking activity in each region. In each population, the incoming synaptic weights are multiplied by the presynaptic activities, to produce the total postsynaptic current. In turn, this current is summed to a noise term and passed through a threshold activation function, to produce a binary activity: 0 if the cortical population is silent, and 1 if it is firing. We derived the resting state fMRI signal from each region as the total postsynaptic current of the corresponding excitatory population.

We computed the cortico-cortical anatomical connectivity that we inserted in the model as the number of connections from the entire cortical source region to the unit volume of the cortical target region, estimated from the Allen Mouse Brain connectome [23–25, 50]. The anatomical connectivity was multiplied by a global scaling coefficient (a free parameter determined by best fit which was the same for all the connections in the model), which represents the average synaptic efficacy per unit of anatomical connectivity strength. In keeping with previous investigations, inter-regional E to E connectivity within and across hemispheres was assumed to be symmetric across the sagittal plane [23, 25, 50]. Resting state fMRI BOLD activity in each region resulting from the underlying neural activity was modeled as the total input current to the excitatory population in each region (Fig. 1B). This assumption is supported by the finding that BOLD correlates best with the local field potential (LFP) [51, 52], and that the LFP reflects the synaptic inputs to excitatory neurons [51, 53, 54].

The free parameters of our network model were the global scaling coefficient of the anatomical connectivity (*G*_*E,E*_), the strength of the I to E, E to I and I to I connections (*J*_*E*,*I*_, *J*_*I*,*E*_, *J*_*I*,*I*_, respectively), the membrane potential firing threshold (*V*^thr^), and the standard deviation of the noise sources (σ). We chose the values of these parameters so that they provided the best fit of the time-averaged mean of the fMRI signal in each region and of the subject-averaged Functional Connectivity (FC) between all pairs of regions that were computed from the empirical mouse resting-state fMRI data obtained in our laboratory (see Methods). We report the best-fit values of these parameters in Table S1.

To evaluate how closely our model reproduced empirical resting state fMRI time series, we first compared time-averaged fMRI activity estimated by the model (Fig. 2A; for comparison we report also the model’s average spiking activity of the excitatory populations in Fig. 2C) with empirical fMRI data (Fig. 2B). The topography of fMRI activity of the model reproduced that of the empirical data remarkably well (ρ = 0.85, p < 10^-5^). The model also closely reproduced two hallmark features of resting state fMRI organization in the mouse brain, namely a highly synchronous resting state fMRI activity within the Default Mode Network (DMN) and Lateral Cortical Network (LCN) [45], and the presence of a robust inter-hemispheric fMRI coupling in homotopic regions, exemplified by the symmetry of the time-averaged fMRI activity between left and right hemispheres [55].

**Figure 2:**
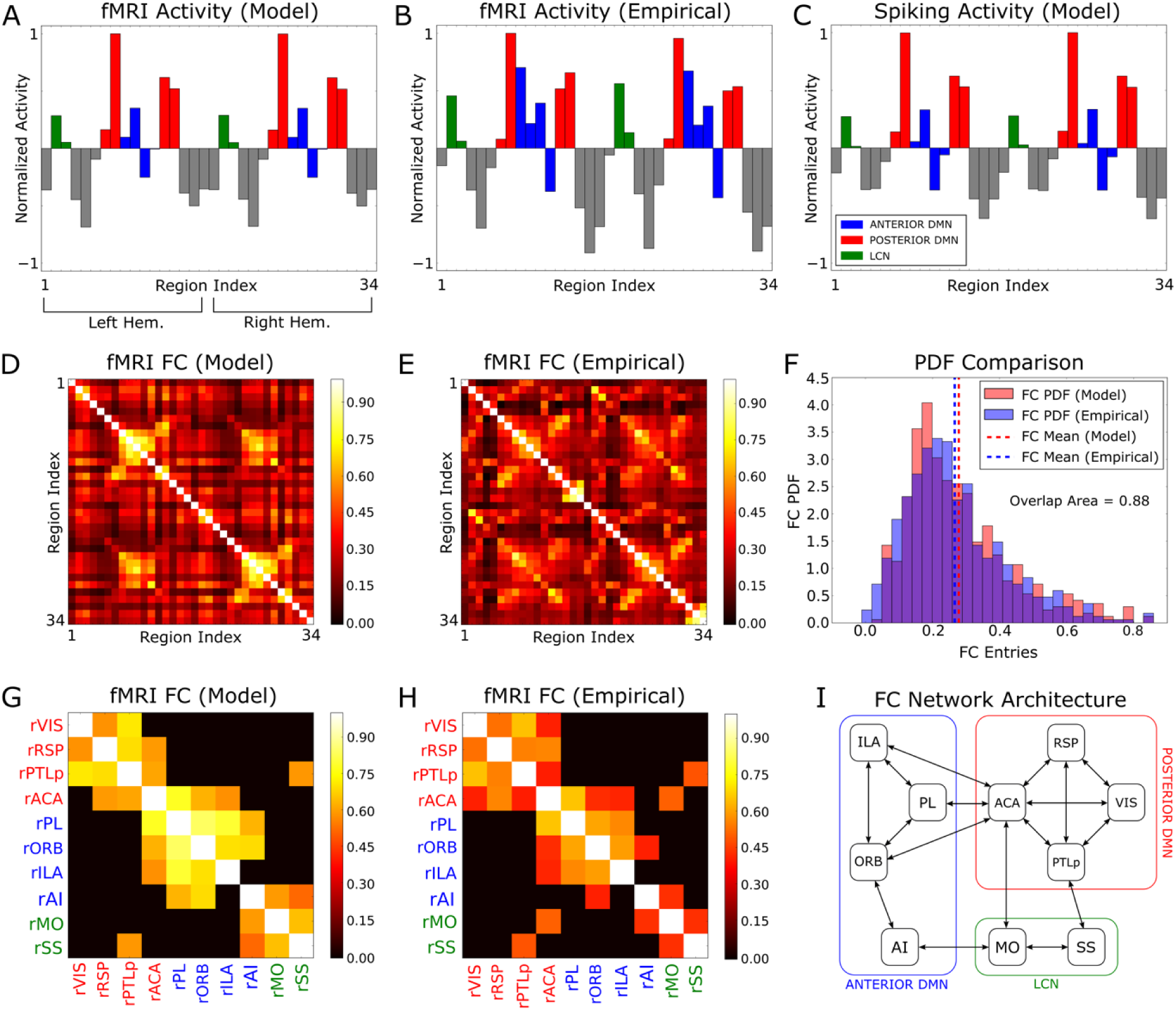
Whole brain modelling recapitulates static fMRI organization of the mouse cortex. **A)** Relative across-region distribution of the time-averaged fMRI activity predicted by the model. For this calculation, the time-averaged fMRI values were normalized between -1 and +1 as explained in Methods. **B)** As in A, but obtained from the empirical mouse fMRI data. **C)** Relative across-region distribution of the time-averaged spiking activity of the model’s excitatory populations. **D)** Static FC matrix of model fMRI time series. **E)** Static FC matrix of the empirical mouse fMRI data. **F)** Comparison between the probability distributions of the values of the entries of the FC matrices, obtained from panels D, E. **G)** Model FC matrix between the regions in the right hemisphere with the highest average resting state mean activity. To highlight the most functionally connected regions, the FC matrix was thresholded to 0.6 of the maximum off-diagonal entry. **H)** As in G, but for the empirical FC matrix. In panels G, H, for simplicity, we showed only the reduced FC matrices in the right hemisphere, while the same results are obtained for the left hemisphere. **I)** Architecture of the functional subnetwork shown in panel H.

We next considered how well our model reproduced static fMRI functional connectivity (FC). We found that the topography of the model’s FC (Fig. 2D) resembled well (ρ = 0.55, p < 10^-5^) that of the empirical FC (Fig. 2E). Importantly, modelled fMRI activity closely reconstituted the range of FC values measured in the real data, as the distribution of FC matrix entries (Fig. 2F) was remarkably similar between model and empirical data, with very high overlap of 0.88 between distributions.

To further demonstrate that modelled fMRI activity closely reproduced hallmark features of the mouse resting-state static FC, we focused on the FC between the regions with the highest average resting-state activity (Figs. 2G, H). Our model reproduced three main cortical subnetworks characterized by higher activity, namely the posterior (VIS, RSP, ACA, PTLp) and anterior DMN (PL, ILA, ORB, AI), and motor regions (MO, SS) belonging to the LCN. This division into subnetworks was further confirmed by applying the leading eigenvector method [56] to determine the community structure of the subnetworks (see Methods), resulting in the functional subnetwork architecture schematized in Fig. 2I.

To probe whether the model explained features of the FC that go beyond its similarity with the anatomical connectivity, following [13] we calculated the correlation between the theoretical FC matrix of the model and the empirical FC matrix, partialized on their common anatomical connectivity matrix. We found that the resulting partial correlation values were 0.47 and 0.40 (p < 10^-4^), when evaluated on the upper and lower triangular parts of the matrices, respectively. This finding implies that our model explains genuine functional interactions among regions, beyond what can be simply inferred from their common anatomical matrix.

In sum, our model closely recapitulates key properties of static cortical fMRI activity and functional connectivity in the mouse brain.

### 2.2 Mouse network model explains anti-correlation patterns in fMRI connectivity after global signal regression

It is well documented that, as we also showed in Fig. 2, the static FC between cortical regions computed on time scales of minutes has mostly positive entries [33, 55]. However, when regressing out the fluctuations of global, whole-brain fMRI activity (“global signal”), human and animal fMRI has reported the formation of robust anti-correlations between specific brain regions [57–60]. In the mouse brain, after global signal regression negative correlations emerge between associative and unimodal sensory-motor regions of the DMN and LCN [32]. We hypothesized that large-scale networks could generate global fluctuations of overall mean activity, which would in turn push the time-averaged FC matrix toward positive values. This, in turn could obscure more nuanced short-timescale interactions between regions that depend on the anatomical interactions and that could be revealed after regressing out global fluctuations.

To test this hypothesis, we computed, both in the network model and in the empirical fMRI data, the global-signal-regressed FC matrix. Note that the FC computed in this way captures not only static connectivity but also frame-wise interactions between local and global activity.

In Fig. 3 we report the FC matrices obtained from the model and empirical fMRI time series, after global signal regression [58]. A strong similarity between these two matrices was apparent when plotting results across all pairs of regions (Fig. 3A vs Fig. 3B, ρ = 0.57, p < 10^-5^) or when zoomed in across the subset of 10 regions with stronger mean activity, including both intra- and inter-hemispheric FC (Figs. 3D-G). Both the empirical and model FC matrices reveal prominent positive correlations between DMN regions ACA and RSP, and negative FC with the LCN region SS. The global-signal-regressed FC of the model and the data were also similar in terms of the distribution of values across entries (Fig. 3C, overlap of 0.82). It is important to note that in our fitting procedure we did not attempt to maximize the similarity of the FC matrices obtained after the global signal regression. For this reason, the strong correspondence between empirical and model global-signal-regressed FC matrices does not trivially reflect the similarity of the FC obtained without global signal regression (Figs. 2D, E), but instead reflects the model’s ability to correctly capture the relevant interactions between global and local activity fluctuations, even when not explicitly fitted to do so.

**Figure 3:**
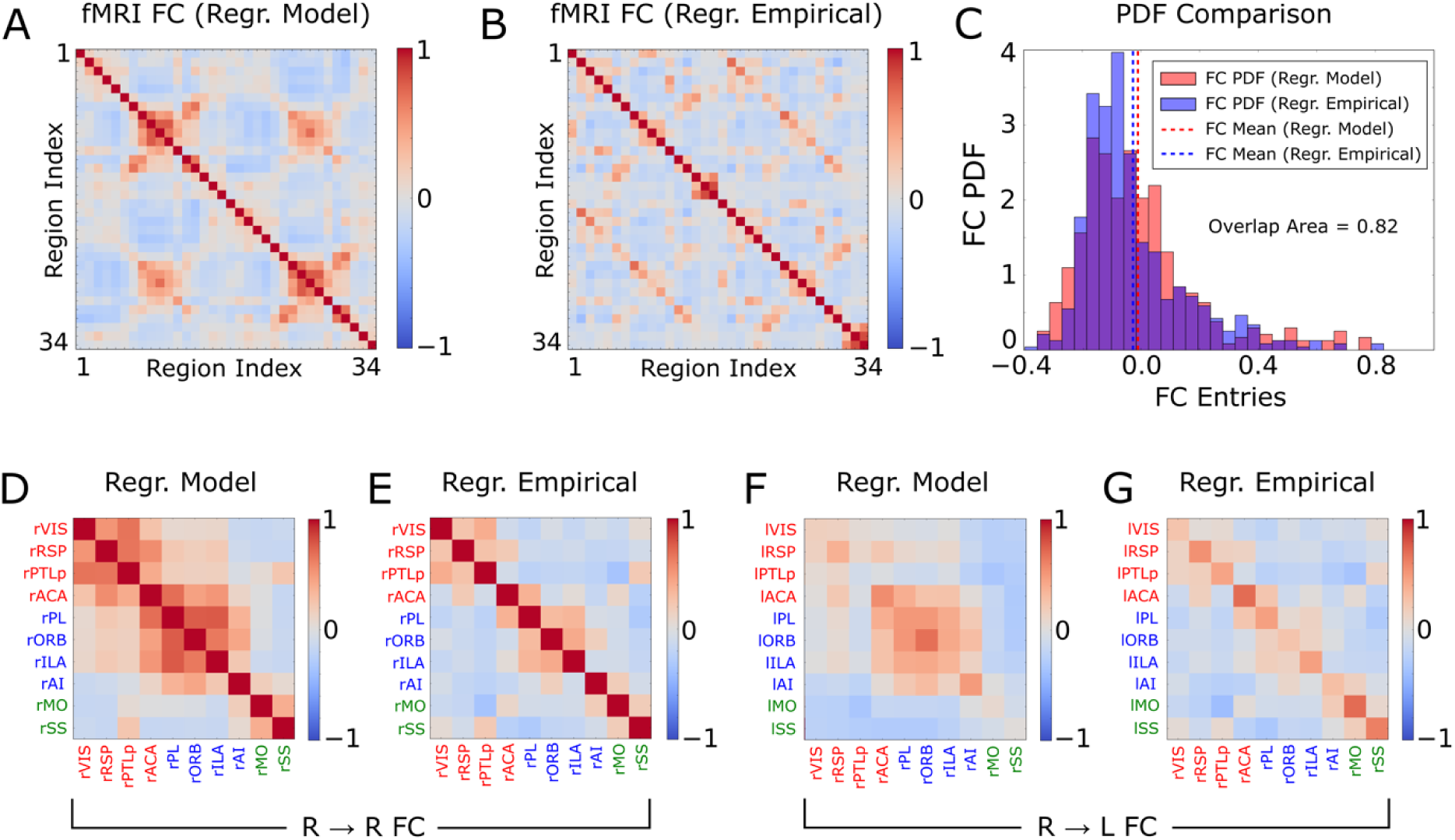
Model and empirical functional connectivity after global signal regression. **A)** Static FC matrix obtained from the model fMRI activity. **B)** As in A, but for the empirical fMRI activity. **C)** Comparison between the probability distributions of the values of the entries of the FC matrices, obtained from panels A and B. **D-E)** Intra-hemispheric FC matrices obtained by restricting, respectively, panels A and B to the 10 regions with strongest BOLD signals in the right hemisphere (similar results are obtained for the L to L connections, not shown). **F-G)** As in D-E, but for the R to L inter-hemispheric connections (similar results are obtained for the L to R connections, not shown).

### 2.3 The network model generates attractor dynamics that explains properties of empirical fMRI co-activation patterns

Previous studies in humans and animals showed that resting state fMRI activity is characterized by a rich temporal organization [37, 61]. One prominent feature of this temporal structure is a dynamic reconfiguration, on the timescale of seconds, into transient brain-wide network states, known as fMRI co-activation patterns (CAPs). CAPs have been consistently found in both humans [35, 62] and mice [30, 31] by clustering frame-by-frame whole brain fMRI activity into a small number of clusters that are robustly detected across datasets, individuals or conditions. However, theoretical conceptualization of how CAPs emerge and are shaped by the concerted interaction of brain regions at rest remains debated. To address these questions, we used our network model to relate CAPs measured from empirical fMRI mouse data to the underlying collective dynamics of cortical regions.

It is well known that models of recurrently connected neural networks like the one we employed here exhibit attractor dynamics [48, 49, 63]. For each attractor, its basin of attraction represents the set of initial activity patterns that eventually end up into that attractor in absence of noise (Figs. S1A, B). In the presence of noise, the network dynamics wanders between attractors. We thus hypothesized that attractor dynamics may also exist in empirical mouse fMRI timeseries, and that the emergence and features of CAPs may be related to those of attractors.

To probe this hypothesis, we analyzed the dynamics of our model numerically (see Ref. [46] and Methods). In the absence of noise, our network converged either to a fixed point where the spiking activity remains constant over time (similarly to the Hopfield network [63]), or to a periodic oscillatory sequence of activity patterns. In the presence of noise, the spiking activity of our whole-brain model featured transitions, over a few-seconds-timescale (Fig. S3A), between attractors. Because each spiking activity attractor state of the network model corresponded to a different simulated fMRI pattern, the attractor dynamics was also visible on the simulated fMRI activity (see Methods).

We found that our network model with the best-fit parameters had 31 stationary states and several thousand oscillatory states (Table S2). To illustrate the topography of the stationary states, for each of the 31 stationary attractors we computed the mean z-score vectors of the model’s fMRI. Interestingly, our model predicted that 7 out of 31 stationary attractors (1, 7, 11, 17, 23, 30, 31, plotted in Figs. 4A and 5A) were homotopic (i.e., with mirror-symmetric activity across the sagittal plane). The remaining stationary attractors were non-homotopic, and occurred in pairs with spatially opposed configurations ((2, 6), (3, 14), (4, 20), (5, 25), (8, 15), (9, 21), (10, 26), (12, 16), (13, 27), (18, 22), (19, 28), (24, 29), plotted in Figs. 4B and 5B).

**Figure 4:**
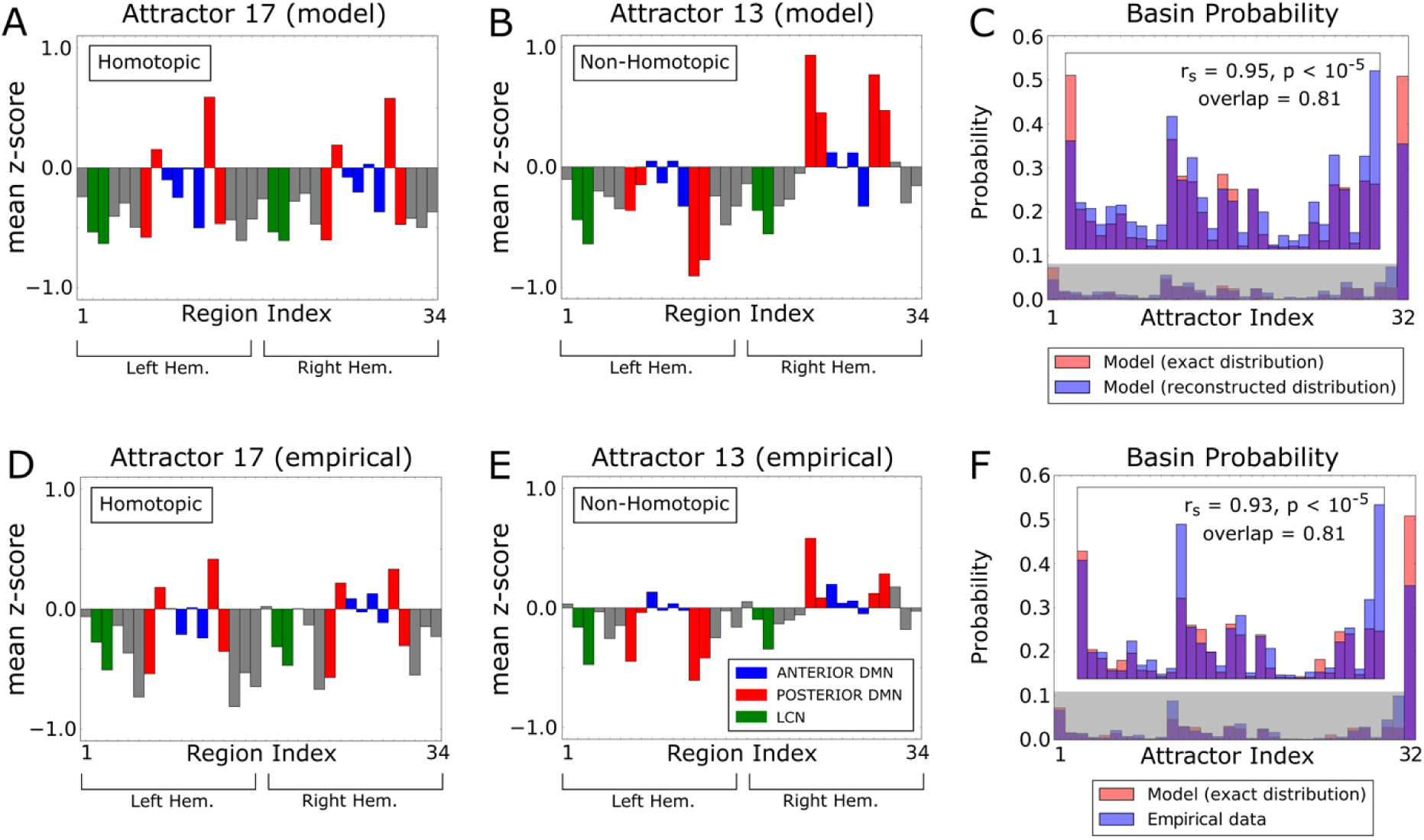
Attractor dynamics of the mouse cortex reconstructed by the mapping algorithm. **A)** Mean z-score vector of model fMRI activity classified into the basin of an attractor (#17) by the mapping algorithm. This attractor is homotopic. **B)** As in A, but for the basin of attractor #13. This attractor is non-homotopic. **C)** Probability distribution of the basins of attraction, calculated numerically from the spiking activity of the model (red bars), and reconstructed by the mapping algorithm when applied to the model fMRI activity (blue bars). The figure inset shows a zoom of the probability distribution of the stationary attractors in the shaded grey area. **D-E)** As in A-B, but in this case the z-scores have been averaged over the empirical fMRI states that the mapping algorithm associated to the basins of attractors 17 and 13, respectively. **F)** As in C, but now the blue bars represent the probability distribution of the basins of attraction reconstructed from the empirical fMRI activity.

**Figure 5:**
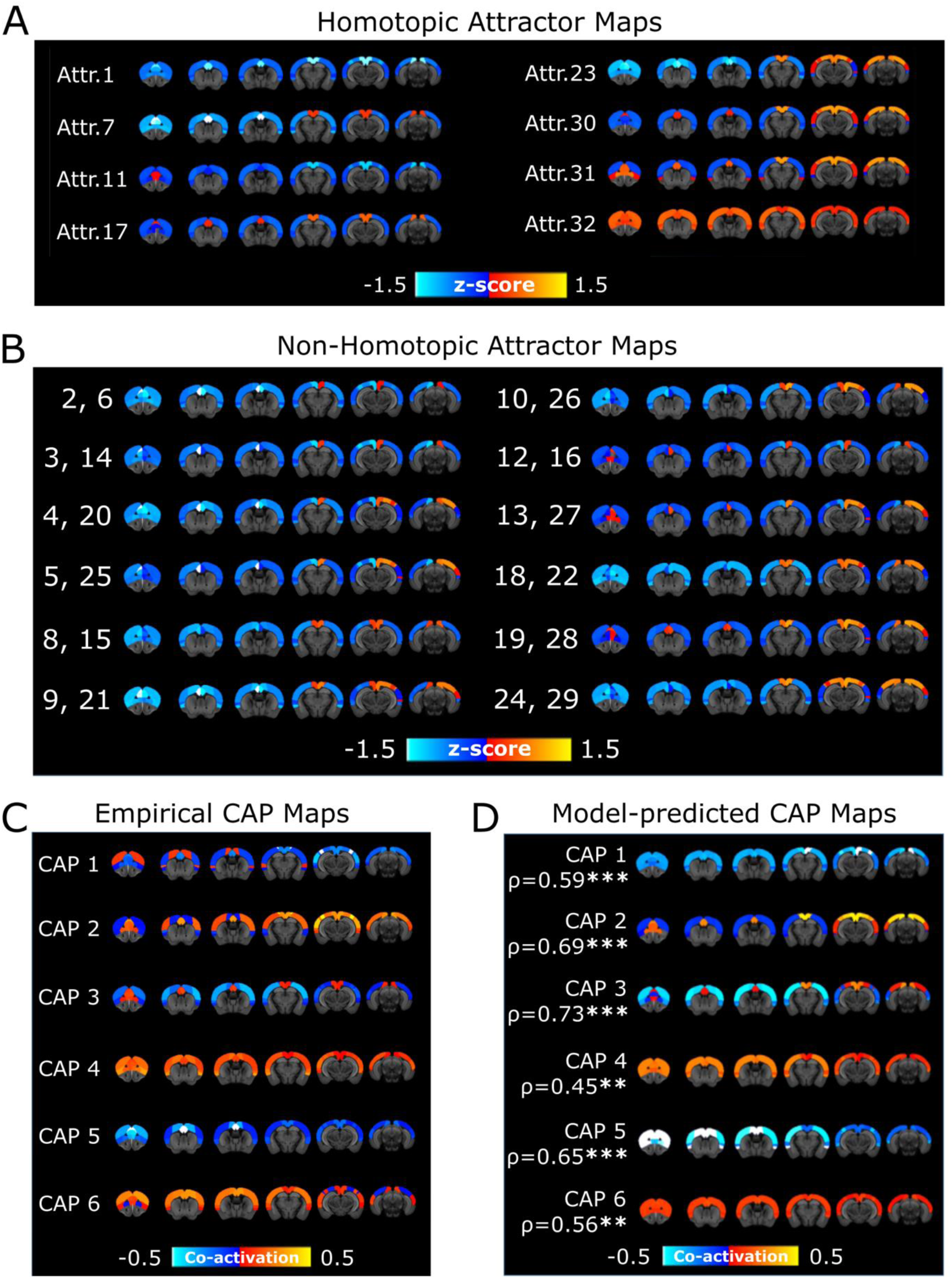
fMRI co-activation patterns and attractors in the mouse cortex A-B) Topography of the homotopic and non-homotopic stationary attractors predicted by the model. **C)** Topography of the CAPs obtained from the empirical fMRI time series. Red-yellow indicates co-activation, while blue indicates co-deactivation. **D)** Topography of the CAPs reconstructed from the model attractors and their Pearson’s correlation with the corresponding empirical CAPs.

The formation of non-homotopic attractors can be explained as arising as a consequence of the non-linear phenomenon, known both in Theoretical Physics and neural network theory, of Spontaneous Symmetry Breaking [64–66] (see section S1 for an illustration in a simple neural network model). Moreover, the fact that the non-homotopic attractors occurred in pairs of spatially opposed configurations suggests that the presence of strong inter-hemispheric asymmetry in frame-by-frame activity may be masked by the left-right inter-hemispheric symmetry of fMRI activity when averaging or pooling data over long timescales. We plotted the probability that each attractor occurred in the model in Fig. 4C (red bars), showing that some stationary attractors (i.e. 1, 11, 12, 13, 16, 17, 19, 27, 28, 30, 31) were far more likely than others.

To probe if empirical fMRI time series showed signatures of dynamical attractors similar to those predicted by our model, we mapped algorithmically each point of a fMRI time series into the basins of the attractors predicted by the model (see Methods and Figs. S1C, D). By averaging over the fMRI time points assigned to each basin of attraction, we could estimate the typical spatial fMRI activation pattern of each putative basin. By counting the number of time points assigned to each basin of attraction, the algorithm next estimated the probability with which each basin is visited over time. Since the probability of each oscillation attractor basin is typically very low (< 10^-5^ on average), it would be unrealistic to reconstruct them with a limited number of empirical samples. For this reason, we mapped only the basins of the individual stationary attractors, and collapsed the basins of the oscillatory attractors into a single macroscopic basin. By comparing the topography of the basins of attraction and the occupation time predicted by the model with those estimated from the fMRI time series, we could quantify if the empirical fMRI time series were compatible with the given attractor dynamics.

We validated the attractor mapping algorithm on the fMRI time series simulated from our whole-cortex network model with the best-fit parameters. The probability of occupancy of the basins reconstructed algorithmically from the simulated time series matched very well the one computed by the theoretical analysis of the model (Fig. 4C, r_s_ = 0.95, p < 10^-5^, overlap area 0.81).

To further validate the mapping algorithm on the fMRI time series simulated from our model, we varied systematically the degrees of inter-hemispheric connectivity, sparseness of the anatomical connections, and of local E-I balance around the best-fit values (see Sections 2.4 and S2 for more details of how the parameter manipulations were done). Then, we applied the attractor mapping algorithm embedded with the attractor structure of the best-fit model to the simulated time series. These manipulations change the number and structure of attractors. As expected, see Figs. S2A-C, the time series with the highest match with the embedded attractor structure were those generated by the model with the best fit parameters (r_s_ = 0.95, p < 10^-5^, mean correlation across all attractors ρ = 0.94, p < 10^-5^), while a different set of model parameter values produced a weaker match.

To investigate whether the empirical time series of fMRI activity showed a temporal structure compatible with the attractor dynamics of the best-fit network model, we next applied the attractor mapping algorithm embedded with the attractor structure of the best-fit model to the empirical fMRI data. The mapping algorithm reconstructed very well the topography of each basin of attraction (mean correlation across all attractors ρ = 0.78, p < 10^-3^, see Figs. S3B, and compare Figs. 4A, D with Figs. 4B, E, respectively). The probability of occupancy of the basins of attraction estimated from empirical data also matched remarkably well that obtained from the theoretical analysis of the best-fit model (r_s_= 0.93, p < 10^-5^, overlap area 0.81, Fig. 4F). We next tested the attractor mapping on empirical data by varying systematically the degrees of inter-hemispheric connectivity, sparseness of the anatomical connections, and of local E-I balance around the best-fit values. Computing this match with a different set of model parameter values produced a weaker match between fMRI and model attractors (Figs. S2D-F). Thus, the frame-by-frame dynamics of the empirical fMRI time series is compatible with attractor dynamics of the specific form generated by the best-fit model, despite the parameters of our model not being optimized to fit frame-by-frame dynamics.

We next asked whether the model’s attractor dynamics could explain the emergence of CAPs in empirical fMRI data. For each time frame of empirical mouse fMRI activity, we compared the CAP it was assigned to [30], to the cortical model’s attractor it was mapped into by the attractor mapping algorithm. Following previous work [30, 31, 37], CAPs were obtained by clustering all the whole-brain empirical fMRI time series conservatively into 6 different clusters, each representing a recurring pattern of fMRI co-activation that is robustly conserved across datasets and individuals. This revealed three distinct CAPs each with a complementary anti-CAP state, i.e. a mirror motif exhibiting a strongly negative spatial similarity to its corresponding CAP (Fig. 5C). Each imaging time frame was mapped to a single CAP and a single basin of attraction. Given that there are more model attractors than empirically determined CAPs, each of the 6 CAPs could potentially capture the contribution of several attractors. We thus reconstructed each CAP from the model’s attractor dynamics as a sum of the spatial activation of each basin of attraction, weighted by the empirical probability that a data point assigned to a CAP was assigned to any given basin of attraction (Fig. S3C). We found a good correspondence between empirically determined and model-attractor-predicted CAPs (Fig. 5D, 0.45 ≤ ρ ≤ 0.73, p < 10^-2^, all CAPs), suggesting that the attractor dynamics of our model is predictive of the topography of empirical CAPs. Importantly, this correspondence between the frame-by-frame dynamics of the model and that of the real data was obtained despite the model’s parameters being not fitted to replicate empirical CAPs.

The match between the topography of empirical and model CAPs was maximal when using model parameters determined by best fit of the static fMRI activity and connectivity. Computing this match with a different set of model parameter values produced a weaker match (Figs. S2G-I). Since model parameters different than the best-fit ones produced a different set of attractors, this result further shows that the frame-by-frame dynamics of the empirical fMRI time series is compatible with attractor dynamics of the specific form generated by the best-fit model.

The CAPs predicted by the network model are also broadly consistent with the FC obtained after global signal regression. The antagonistic interaction between DMN and LCN network hubs (Fig. 3) is also observed in the transient, opposite co-activation of empirical CAPs (Fig. 5C), which were well predicted by the reconstructed CAPs through the model attractor dynamics (Fig. 5D, CAPs 2 and 3). Other CAPs with more global cortical synchrony (CAPs 4 and 5) show transient co-activations between regions from both DMN and LCN, corroborating the contribution of CAPs to FC structure as a weighted sum of transient correlations that add up to the observed FC matrices. We also noted that CAPs 2, 4 and 6 (called “anti-CAPs” in [30] as they look like reversed activation patterns with respect to CAPs 1,3 5 and are observed in opposite phases of the global signal fluctuations with respect to their accompanying CAPs 1, 3 and 5) have a stronger contribution from oscillatory attractors than the corresponding CAPs 1, 3, 5 (Fig. S3C). This suggests that the antagonist nature of CAPs and anti-CAPs may be partly related to an antagonism of transitions between stationary and oscillatory attractor types.

Together, these findings suggest that CAPs are, at least in part, an observable manifestation of a whole-cortex attractor dynamics of the type observed in our network model.

### 2.4 Inter-hemispheric coupling, anatomical connectivity sparseness and local excitation-inhibition balance shape model cortical attractors

We demonstrated above that interactions between anatomically connected cortical regions drive attractor dynamics and network states. In this section, we investigate how these dynamics depend on three key properties of cortico-cortical connectivity: inter-hemispheric connection strength, connection sparseness, and the balance between excitatory and inhibitory populations. This dual-purpose analysis aims to deepen our theoretical understanding of how attractors emerge from connectivity patterns, and to determine whether specific features of mouse brain connectivity uniquely support rich attractor dynamics.

First, we modified the inter-hemispheric connectivity by means of a global inter-hemispheric scaling coefficient *W*, which multiplies the weights of all the inter-hemispheric anatomical connections. *W* = 0 corresponds to the case when the two hemispheres are anatomically disconnected, while *W* = 1 corresponds to the original empirically measured values for the connectome.

The dependence of the number of attractors on the inter-hemispheric connectivity scaling parameter *W* is shown in Fig. 6A. The total number of attractors decreased for high values of *W* which correspond to abnormally strong inter-hemispheric connectivity (*W* ≫ 1). An intuitive explanation for this is that excessively strong inter-hemispheric connectivity would increase the overall input current to each cortical region, thereby increasing their firing probability. As our model has stochastic noise, this results in the formation of a single stationary attractor, where all the regions fire synchronously.

**Figure 6:**
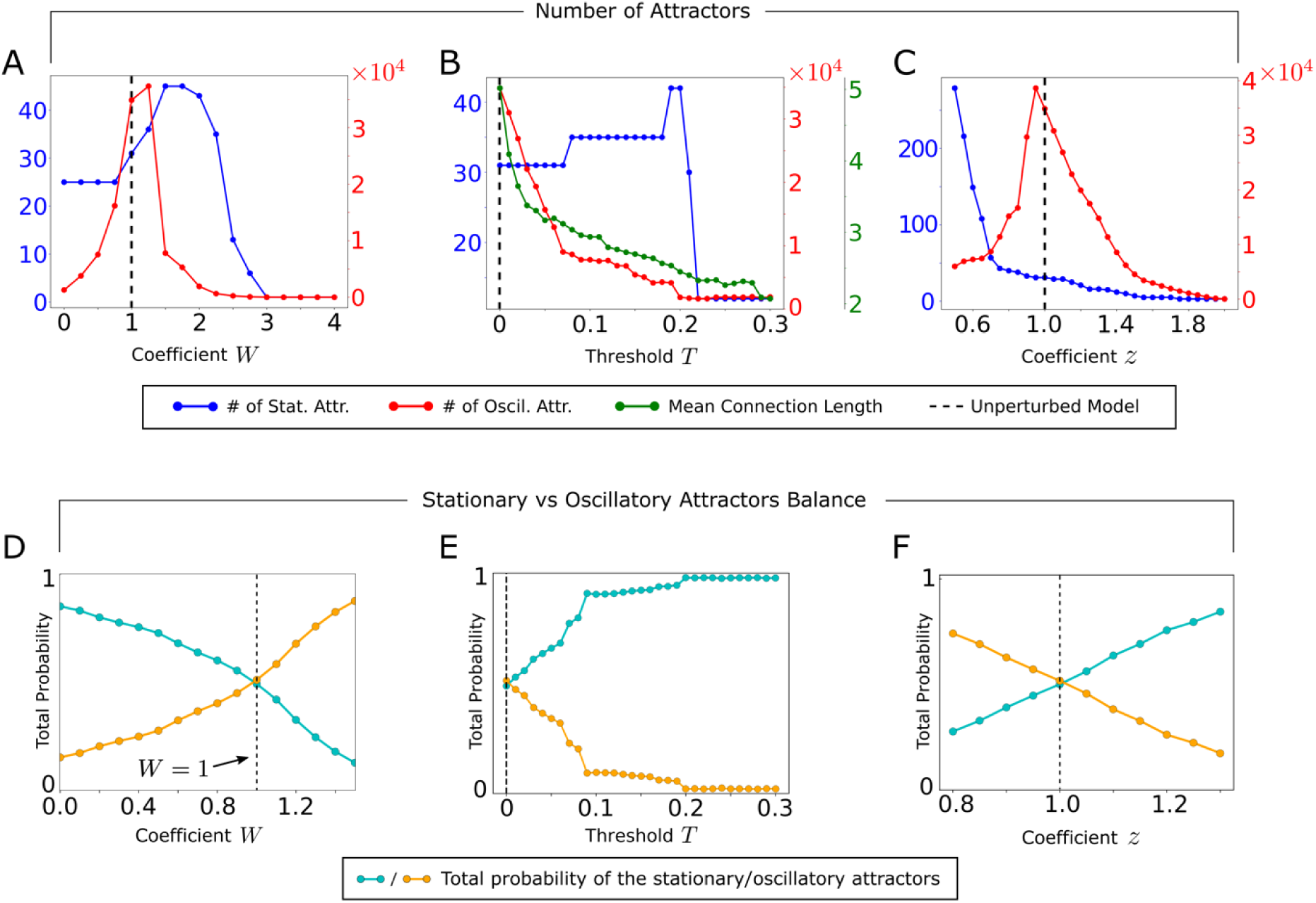
Relationship between attractors and features of anatomical connectivity. **A)** Number of stationary and oscillatory attractors as function of the global scaling coefficient *W* that multiplies the inter-hemispheric synaptic weights. The total number of attractors exhibits a peak for the value corresponding to the real mouse connectome (*W*∼1). **B)** As in A, but as function of the threshold value *T* used for sparsifying the anatomical connectivity. **C)** As in A, B, but as function of the scaling coefficient *z* that multiplies the E to I and I to E couplings between populations. **D)** Probability to observe the activity pattern in the basins of attraction of the stationary states (cyan curve), or in those of the neural oscillations (orange curve), when varying *W*. Note that for *W* = 1, the model exhibits an approximate balance between the two probabilities. **E)** As in D, but when varying the threshold value *T*. **F)** As in D, E, but when varying the scaling coefficient *z*.

The total number of attractors also decreased when we used inter-hemispheric connectivity values smaller than the one of real data (*W* < 1). In this case inter-hemispheric input adds only marginal variability to the total input to each region. As a result, the network accesses fewer attractors. The number of attractors peaks for intermediate values of *W*. Specifically, the number of oscillatory attractors, which dominates the total number of attractors, has a local peak for the original model of the mouse brain (*W* = 1), see Fig. 6A.

Moreover, the model predicts that the total occupation probability of the basins of attraction of stationary (respectively oscillatory) states decreases (respectively increases) with *W*, and the real-connectivity value corresponds to approximate balance between the time that the network spends in stationary vs oscillatory attractors (Fig. 6D). These results suggest that the mouse brain operates at an optimal point for attractor richness.

We next investigated the role of the sparseness of the anatomical connections in creating attractors. We sparsified the anatomical connections between the excitatory populations by removing all the links below a threshold *T*. The case *T* = 0 corresponds to the original connectome, while larger values of *T* produce sparser versions of the connectome containing only the strongest connections. In Fig. 6B we report how the number of stationary and oscillatory model attractors depends on *T*. The number of stationary attractors was relatively stable until *T* = 0.2 and then dropped dramatically, suggesting that creation of stationary attractors requires necessarily a skeleton of stronger connections. The number of oscillatory attractors declined strongly but less abruptly with increasing *T*, with a good number of oscillatory attractors still surviving for *T* > 0.2. This suggests that, although stronger connections create oscillatory attractors, weaker anatomical connections can too create oscillatory attractors (see Supplemental Information, section S3 for more details). Note that the weakest links are also the longest, because the mean length of the connections in the network decreases with *T* (Fig. 6B). Thus, our results support recent suggestions [67] that the longest connections add to network dynamics. Moreover, the model predicts that the total occupation time of the basins of attraction of stationary (respectively oscillatory) states increases (respectively decreases) with *T*, and the real-connectivity sparsity corresponds to approximate balance between the time that the network spends in stationary vs oscillatory attractors (Fig. 6E).

Finally, we investigated the role of the coupling strength between excitatory and inhibitory populations. We modified this coupling strength by means of a global scaling coefficient *z*, which multiplies the E to I and I to E connection weights. The value *z* = 0 corresponds to when the excitatory and inhibitory populations are disconnected, while *z* = 1 corresponds to the original network model with the best-fit couplings. In Fig. 6C we show how the number of attractors in the model is affected by the scaling coefficient *z*. The number of oscillatory attractors, and therefore also the total number of attractors in the network, peaks for values of excitation-inhibition balance equal to those of the best-fit model (*z* = 1). This is because *z* regulates the effective strength of the recurrent E to I and I to E connections that sustain oscillatory dynamics in neural networks [68]. The number of stationary attractors decreases with *z*, because the local I to E synaptic current in each cortical region has a destabilizing effect on the stationary states, which becomes stronger when increasing *z* (see Eq. (S4)). Moreover, the model predicts that the total occupation probability of the basins of attraction of stationary (respectively oscillatory) states increases (respectively decreases) with *z*, and the best-fit *z* = 1 value corresponds to approximate balance between the time that the network spends in oscillatory vs stationary attractors (Fig. 6F). By perturbing the local E-I couplings, we also predicted that the balance is optimal for encoding external stimuli (see Section S5).

### 2.5 Non-homotopic attractor dynamics due to inter-hemispheric anatomical coupling

We next considered the inter-hemispheric topography (that is, the differences between the left and right hemispheres) of the attractor dynamics of the model. Given that excitatory inter-hemispheric coupling should intuitively synchronize homologous regions across the hemispheres, we investigated how the strength of those couplings affects the homotopicity of the attractors. We found (Fig. 7A) that the number of non-homotopic attractors (i.e., with mirror-asymmetric activity across the sagittal plane) was small for large values of *W*, as expected by the simple intuition that strong inter-hemispheric coupling leads to strong inter-hemispheric synchronization. Surprisingly, the number of non-homotopic attractors peaked at intermediate values of inter-hemispheric connection strengths (*W*∼1) rather than at null inter-hemispheric strength (Fig. 7A). This result was confirmed by computing (Fig. 7B), as function of *W*, the inter-hemispheric Hamming distance between spiking activity of homologous excitatory populations across the hemispheres. The latter is a measure of non-homotopicity that does not rely on computing attractors.

**Figure 7:**
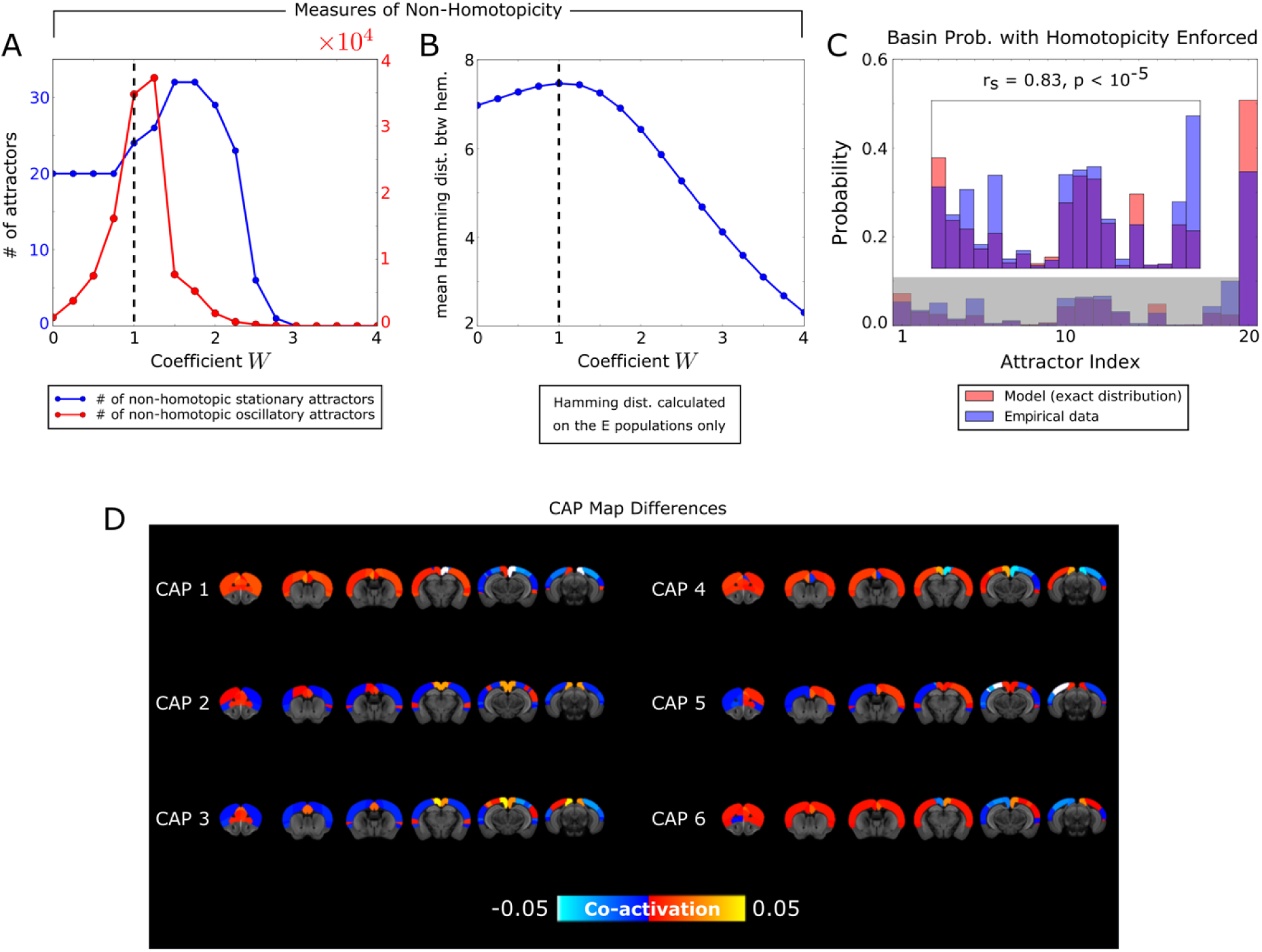
Model study of non-homotopic patterns of fMRI activity. **A)** Number of non-homotopic attractors in the network model, as function of the global scaling coefficient *W*. **B)** Mean Hamming distance between the spiking activity patterns of the excitatory populations in the L/R hemispheres. **C)** Probability distribution of the basins of attraction, calculated numerically from the spiking activity of the model (red bars), and reconstructed by the mapping algorithm when applied to the empirical fMRI signals (blue bars), using the attractor structure with homotopicity enforced. Compare this result with Fig. 4F (obtained for mapping the empirical data into the attractor structure of the original model without enforcing homotopicity). **D)** Difference between the CAPs of the model with non-homotopic attractors, and the corresponding CAPs as obtained from the model with enforced homotopicity of attractors.

Thus, although our best-fit model exhibited dynamics that was homotopic when averaged over long-time scales of minutes, it did instead predict, for realistic values of inter-hemispheric connectivity (*W* ∼1), the presence of non-homotopic stationary and oscillatory attractors at the time scale of seconds. This suggests that intrinsic fMRI dynamics in the mouse brain may feature significant moments of non-homotopic activity arising from attractor dynamics, even if the overall static average remains homotopic.

To assess whether the non-homotopicity in the model attractors had a counterpart in the empirical fMRI data, we used our algorithm to map the data into either the attractor structure of the original model (as done in Fig. 4) or to an attractor structure obtained by artificially enforcing homotopicity (see Methods). We found that mapping the empirical data onto homotopic model attractors predicted attractor occupancy probability less accurately than mapping it onto the original model with non-homotopic attractors (r_s_ = 0.831 ± 0.004 vs r_s_ = 0.933 ± 0.004, p < 10^-5^, see Fig. 7C). This suggests that the non-homotopicity of the attractors space is important to describe the frame-by-frame dynamics of the empirical data.

The importance of non-homotopic frame-by-frame dynamics is also reflected in the reconstruction of the topography of empirical CAPs from the attractors. In Fig. 7D we report the differences between the CAPs of the original model with non-homotopic attractors, and the corresponding CAPs as obtained from the model with enforced homotopic attractor structure. CAP 1 and CAP 6 exhibited significant decrease in the correlation with the corresponding empirical CAPs when using homotopic attractors, while the remaining CAPs did not exhibit significant variation. Specifically, the correlation for CAP 1 decreased from 0.588 ± 0.007 to 0.562 ± 0.005 (p < 10^-5^), while for CAP 6 it decreased from 0.561 ± 0.007 to 0.512 ± 0.007 (p < 10^-5^). Thus, the non-homotopicity of model attractors gives some contribution to explaining some of the features of the empirical data that is lost when artificially making them homotopic. The formation of non-homotopic activity in the presence of inter-hemispheric excitatory coupling can be interpreted as a form of Spontaneous Symmetry Breaking [64]. We report in Supplemental Information, section S1, an explanation of this mechanism for neural networks. Together, these results suggest that the inter-hemispheric exchange of information through the corpus callosum and the anterior commissure pathways can create non-homotopicity of activity and possibly support dynamical differences in information processing between hemispheres.

### 2.6 Importance of directionality of connectivity and non-linearities in fMRI neural dynamics

Our network model included two important aspects of biological plausibility not always present in standard large-scale fMRI modelling: directed anatomical connections and threshold-based non-linear neural dynamics. To test the relative contribution of these features to fMRI activity, we compared our whole-brain model to two simpler models obtained either by symmetrizing the anatomical connections between the excitatory populations (thus making the connectivity un-directed), or by linearizing the network equations. In both cases, we derived a new set of best-fit parameters (Table S1), following the same fitting procedure that we used for the original model (Methods).

We first considered how the models reproduced static properties of the empirical fMRI data. The Pearson’s correlation between the static across-region distribution of the fMRI signals obtained from data and the models (Fig. 8A) was similar across different models but was higher for our model (p < 10^-5^, two-sample Welch’s t-test). Moreover, our model was considerably better than the linear one at reproducing the topography of the static FC matrix (Fig. 8A), although it did not outperform the undirected model. However, unlike our model, the undirected and the linear models failed to accurately reproduce the distribution of values of the empirical fMRI static FC (Fig. 8B). Thus, directionality of anatomical connectivity and neural non-linearities help reproduce the static organization of the empirical fMRI data.

**Figure 8:**
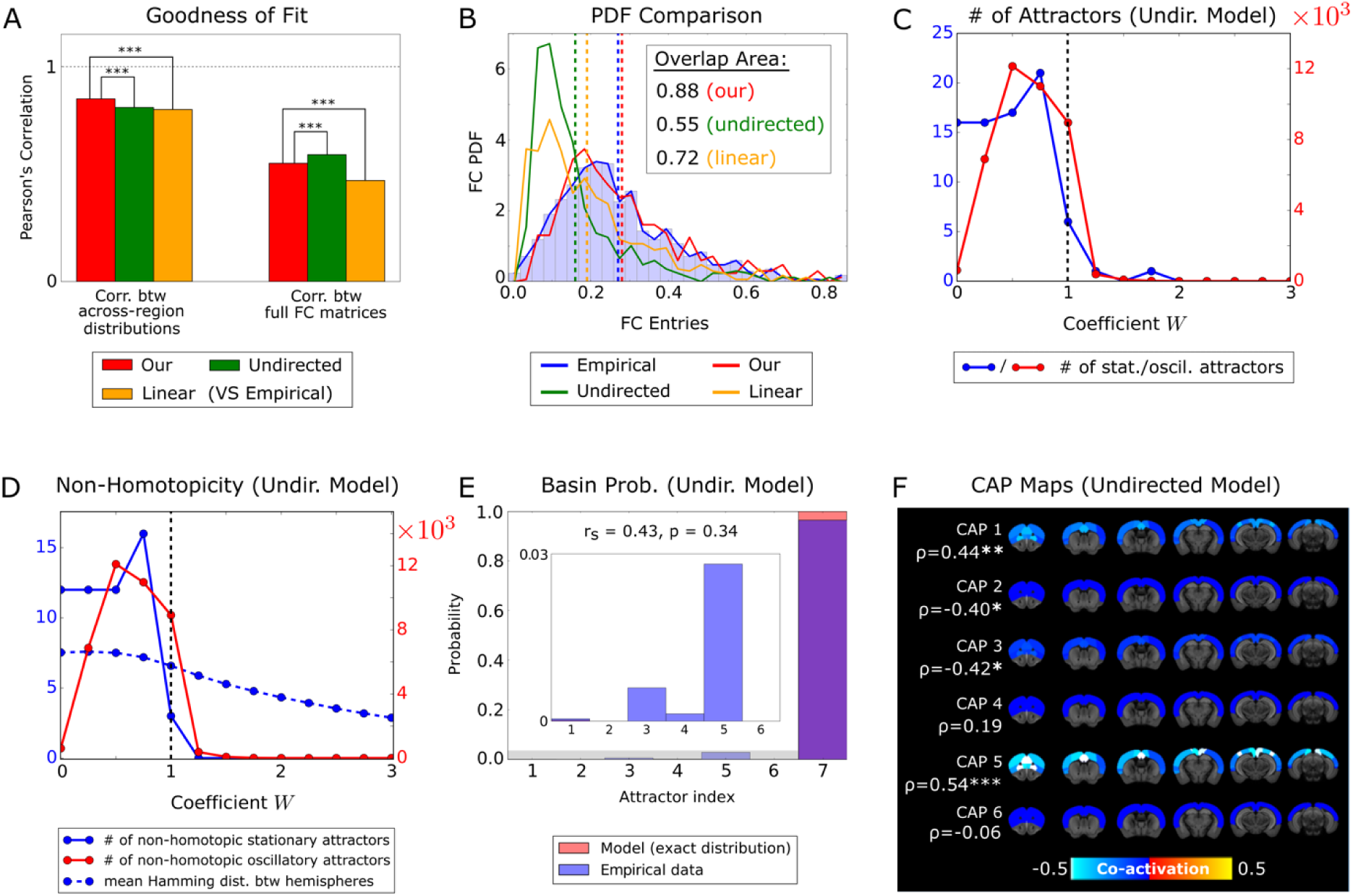
Comparison with undirected and linear models. **A)** Pearson’s correlation between the across-region mean fMRI activity (left) and the FC matrix (right) of the empirical data, and the corresponding statistics calculated for three network models: our full model with directed connectivity and non-linear neural dynamics (red), an undirected model with non-linear dynamics but undirected anatomical connections (green), and a linear model with directed anatomical connections but linear activation function (orange). Statistical comparisons were performed by running, for the three network models, 100 groups of 100,000 repetitions each, and then by using a two-sample Welch’s t-test to compare these distributions. **B)** Comparison between the probability distributions of the values of the entries of the FC matrices. Note that, unlike our model, the undirected and linear models do not fit well the PDF of the empirical datasets. Color coding as in panel A. **C)** Number of stationary and oscillatory attractors of the undirected model, as a function of the global scaling coefficient *W*. Note that the linear model has only one (stationary) attractor. **D)** Measures of inter-hemispheric non-homotopicity for the spiking activity patterns of the undirected model. **E)** Probability distribution of the basins of attraction, calculated numerically from the undirected model (red bars), and reconstructed by the mapping algorithm when applied to the empirical fMRI signals (blue bars). The figure inset shows a zoom of the probability distribution of the stationary attractors in the shaded grey area. **F)** CAPs reconstructed from the fMRI attractors of the undirected model, and their Pearson’s correlation with the corresponding empirical CAPs of Fig. 5C.

We next considered how different models accounted for the empirical fMRI frame-by-frame dynamics. We found a major reduction in the number of attractors generated by the undirected and the linear model compared to our model (Fig. 8C). The undirected model generated much fewer attractors than the original model. For *W* = 1, we found only 6 stationary states in the undirected model, compared to 31 stationary states in our model (Tables S2 and S3), and 8,958 rather than 34,877 oscillations. Moreover, while our model exhibited the largest number of attractors at *W*∼1 (Fig. 6A), for the undirected model that number peaked at *W* = 0.5 (Fig. 8C). Thus, the undirected model exhibits poorer attractor dynamics, which would not be maximized by values of inter-hemispheric anatomical connection strength equal to those of the mouse brain. Similarly, indices of non-homotopic activity (i.e. diverging activity across homologous regions in the two hemispheres) like those considered in Figs. 7A, B did not peak (Fig. 8D) at *W*∼1 for the undirected model, as observed in our model (Fig. 7).

Importantly, we found that the empirical fMRI time series did not map well into the attractors of the undirected model. The Spearman’s correlation between the probability of occupancy of the basins of attraction predicted by the model and the distribution reconstructed by the mapping algorithm was not significant (rs = 0.43, p = 0.34, Fig. 8E), and the undirected model was particularly bad at describing the probability of stationary attractors (inset of Fig. 8E). The impoverishment of attractor dynamics in the undirected model led also to a much poorer reconstruction of the topography of the empirical CAPs from its basins of attraction (cf. Fig. 5C with Fig. 8F). Thus, the undirected model lacks the ability to predict, from attractor dynamics, the features of most empirically observed CAPs.

The attractor dynamics of the linear model was even poorer, as it exhibited only one stationary attractor. Given that there was only one attractor, we did not attempt to map CAPs and attractors in this model.

In sum, the neural non-linearities and the directionality of the anatomical connectivity inserted in our model only marginally increased the accuracy of model predictions of the static organization of empirical fMRI data. However, the non-linearities and the directionality information of the connectome deeply affected the fast network dynamics on the scale of seconds. Specifically, these contributions were required to create richer attractor dynamics in the model, which ultimately allowed it to better explain frame-by-frame empirical CAPs.

## 3 Discussion

Recent years have witnessed a growing interest in modelling and understanding the dynamics of fMRI resting state activity in the mouse brain [18, 38–40]. The mouse has the unique advantage of the availability of a precise measure of whole-brain anatomical connectivity and its directionality [23–25, 45]. Here we contributed to this endeavor by developing a novel whole-cortex model of the resting state activity which adds concepts and predictions to both existing mouse and human whole-brain models [6, 8–10, 13]. Specifically, our model made new predictions about the dynamics of resting mouse cortex in terms of attractor dynamics over fast timescales of seconds, which are compatible with empirical findings on CAP frame-by-frame dynamics. We also manipulated the model’s anatomical connectivity to predict how the structure of cortico-cortical connectivity may generate attractor dynamics and inter-hemispheric non-homotopic activity.

### 3.1 Attractor dynamics may originate from cortico-cortical interactions and may account for the emergence of CAPs

Most whole-brain models simulate the fMRI signals either by modeling neurovascular coupling with a synaptic gating variable [9–11, 13, 69], or by employing neural mass models [6, 12, 14, 15, 70, 71]. Our model instead uses a binary non-linearity for describing neural spiking activity [5, 7], which is known to generate numerically- and analytically-tractable attractor dynamics [46], and models fMRI as postsynaptic currents. Attractor dynamics is a major theoretical feature of recurrently connected neural networks, yet its presence in large-scale brain dynamics has been only seldomly investigated (see e.g. Refs. [7, 13] modelling the human brain). Our work adds to and expands previous attempts to predict attractor dynamics from large scale models in several ways.

First, the model we chose lends itself to an exhaustive coverage of attractor states including stationary and oscillatory attractors. We could map 31 stationary attractors and thousands of oscillatory attractor in resting state dynamics, improving on previous work which mapped no oscillatory attractors in humans or mice [7, 13, 72], or focused on simpler dynamics generated by models of the mouse brain with binarized connectivity [40]. The ability to map extensively attractors was essential to relate the frame-by-frame, fast dynamics of the model to the dynamics of empirical fMRI data on a time scale of seconds. This allowed us to provide predictions and mechanistic hypotheses about possible novel dynamic features such as inter-hemispheric non-homotopicity imputable to attractor dynamics. These predictions extend previous seminal work on attractor dynamics in humans, relating attractor dynamics to time-varying FC [7, 13], and they also extend to the case of resting-state activity the study of attractor dynamics in on whole-cortex models of the mouse during working memory [72].

Second, we investigated the role of the directionality of anatomical fibers, a feature not available to a comparable extent for the human brain, in shaping attractor dynamics. We found that the richness of attractor dynamics and the good match between empirical fMRI dynamics and attractor properties was found only when considering the directionality of the anatomical connectivity of the real mouse brain. Using an undirected version of the anatomical connectivity matrix (conceptually similar to that measured with DTI) led to an impoverishment of the model’s attractor landscape and of the ability to predict real fMRI dynamics. Our determination of the importance for the mouse brain’s attractor dynamics of the directionality of anatomical connections extends previous studies of the role of long-range vs local excitatory connections in producing patterns of internally sustained neuronal activity [72].

Finally, we related attractor dynamics to CAP dynamics found empirically on the time scale of seconds. The possible origin of these CAPs in terms of neural processes has been debated. CAPs have been suggested to reflect widespread cortical changes associated with changes in ascending modulatory input or brain state, or brief, event-like bursts of self-generated activity [35, 41]. Our model provides a possible alternative or additional mechanistic explanation of CAPs. It suggests that CAPs may emerge from and reflect cortico-cortical interactions, and that they can emerge also without the contribution of ascending modulatory inputs. Specifically, our model predicts that CAPs mechanistically arise from attractors, which in turn express and originate from the underlying anatomical cortico-cortical connections and interactions.

Moreover, because attractors represent minima in the energy landscape of the network, our work also suggests that changes in CAPs can be taken as indicators of changes in the energy landscape of the brain, a topic of current active research [43]. It would be important to determine in future theoretical and empirical studies how the interaction between such attractor cortical dynamics and neuromodulatory systems such as the ascending arousal system may shape the energy landscape of the cortex [43].

By enabling us to relate anatomical connectivity, attractors and fMRI CAPs, our formalism might be crucially employed in future studies to make empirically testable predictions about how alterations of the connectome resulting from injury or neurodegenerative disorders may alter frame-by-frame brain dynamics.

### 3.2 Model predictions of fMRI functional connectivity after regressing out global activity

Our model predicted well the empirical FC obtained after regressing out the global signal. While static FC had all positive entries, the regressed FC showed interesting patterns of anti-correlations across regions that were also visible in CAPs and were predicted well by the model. These results contribute to the ongoing debate about the validity and significance of global signal regression in fMRI [73]. While some studies consider the global signal as a confounder or a source of noise (thus advocating its regression to account for in-scanner head motion [74]), other studies provided evidence that the global signal may reflect neural activity by highlighting how it relates to FC topography [75–77], and to CAPs topography and dynamics [30, 31]. Our results support the notion that the global fMRI signal potentially carries important information about how different regions interact to underpin fMRI dynamics.

### 3.3 Model predictions on the relationship between anatomical connectivity and frame-by-frame whole-brain dynamics

We studied theoretically how connectivity strength variations affect the frame-by-frame fMRI dynamics. We obtained several findings. First, values of inter-hemispheric connectivity similar to those found in the real mouse brain maximize the number of attractors. We speculate that the number and variety of possible attractors may impact brain function because it represents the repertoire of brain states that can be reliably and dynamically generated starting from arbitrary initial conditions. Therefore, larger repertoires of attractors may enable a larger repertoire of information processing capabilities [72, 78, 79]. Fitting our model to fMRI activity under different behavioral states could help validating empirically a possible role in brain function of attractor dynamics.

Second, despite our model included an inter-hemispheric homotopic anatomical connectivity matrix (that is, the inter-regional E to E connectivity within and across hemispheres was assumed to be symmetric across the sagittal plane), it predicted the formation of significant differences between the frame-by-frame dynamics of the two hemispheres. Values of inter-hemispheric connectivity strength similar to those of the real mouse brain led to a predicted higher degree of inter-hemispheric non-homotopicity in frame-by-frame cortical activity. This inter-hemispheric non-homotopicity was predicted by the model to happen only over fast timescales (of the order of seconds), but was averaged away when static activity and FC were computed over the several minutes of a whole fMRI time series. This can be conceptualized as spontaneous symmetry breaking, which may aid to create some degree of dynamic dissociation of information processing across hemispheres.

Third, we found that having a good number of stationary attractors necessitates the presence of a backbone of the strongest anatomical connections (which on average have shorter connections). Oscillatory attractor numbers also declined when removing the strongest connections. Since CAPs reflected attractor dynamics in our model, the backbone of strongest anatomical connections may have a more profound influence on CAP formation and structure. A good number of oscillatory attractors, but not of stationary attractors, disappeared when only weak connections were removed in the model (which are also on average the longer-distance ones), corroborating previous proposals that long-distance pathways contribute sizably to whole-brain dynamics [67]. Assuming that larger attractor numbers help information processing, these results are also compatible with other model-based proposals that long-range connections influence information processing in the brain [15].

Finally, our best-fit model parameters were compatible with a local balance in the interactions between excitation and inhibition. This result further supports the notion that aspects of across-region and within-region connectivity combine to strengthen brain computational capabilities.

## Acknowledgements

We thank members of our laboratories for useful discussions and feedback.

## Data Availability Statement

The code and data to reproduce our results is made available and can be found at https://data.mendeley.com/datasets/xscxtshgfx/1.

## 4 Methods

### 4.1 Resting state fMRI data acquisition

The empirical resting state fMRI datasets used in this work were collected from 15 adult male C57Bl6/J mice. These datasets were published and made available previously [30]. To ease reproducibility and development of further modeling work, the fMRI time series of each mouse obtained using the specific parcellation adopted for this paper (see below) are shared again with this paper at https://data.mendeley.com/datasets/xscxtshgfx/1. All in vivo experiments were conducted in accordance with the Italian law (DL 26/214, EU 63/2010, Ministero della Sanità, Roma) and the recommendations in the Guide for the Care and Use of Laboratory Animals of the NIH. Animal research protocols were reviewed and consented by the animal care committee of the Italian Institute of Technology, and Italian Ministry of Health.

Animal preparation, image data acquisition, and image data preprocessing for fMRI data have been described in full detail in [30, 32, 33]. Mice were anaesthetized with isoflurane, intubated and artificially ventilated (2%, surgery). At the end of surgery, isoflurane was discontinued and substituted with halothane (0.75%). Functional data acquisition commenced 45 minutes after isoflurane cessation. The fMRI data were acquired with a 7.0 Tesla scanner (Bruker Biospin, Ettlingen) using a 72 mm birdcage transmit coil, and a four-channel solenoid coil for signal reception. Single-shot EPI time series were acquired using the following parameters: TR/TE 1200/15 ms, flip angle 30°, matrix 100 × 100, field of view 2 × 2 cm^2^, 18 coronal slices, slice thickness 0.50 mm, 500 (n = 15) volumes and a total fMRI acquisition time of 10 minutes, respectively. As in [30], fMRI time series preprocessing included: removal of the first 50 frames (1 minute), despiking, motion correction, and spatial normalization to an in-house mouse brain template with the same native resolution as raw EPI volumes. Head motion traces and the mean ventricular signal (average fMRI time series within a manually-drawn ventricle mask from the template) were regressed out. The resulting images were band-pass filtered using a 0.01 – 0.1 Hz band, and spatially smoothed using a Gaussian kernel of 0.5 mm FWHM.

### 4.2 Mouse anatomical connectivity data and parcellation

The mouse anatomical connectivity data used in this work were derived from a voxel-scale model of the mouse connectome made available by the Allen Brain Institute [23–25]. The mouse anatomical connectome was obtained from imaging-enhanced green fluorescent protein (eGFP)-labeled axonal projections derived by 428 viral microinjection experiments and registered to a common 3D coordinate space. To derive connectivity estimates at the voxel-scale level, the connectivity at each voxel was modeled as a radial basis kernel-weighted average of the projection patterns of nearby injections [24]. Following the procedure outlined in [24], we re-parcellated the voxel scale model according to the highest hierarchical level (i.e. the coarsest parcellation scheme) available in the Allen Mouse Brain Atlas, obtaining 17 cortical regions per hemisphere. We used the connection density (CD) to define the anatomical connectome edges [50]. The same set of 34 cortical nodes was used throughout the work.

### 4.3 CAPs

To identify CAPs in empirical fMRI data we used the mean CAP map templates derived in [30]. Briefly, these CAPs were identified by clustering the concatenated fMRI frames of N = 15 mice using the k-means++ algorithm, with 15 replicates, 500 iterations, and Pearson’s correlation as distance metric. Previous work [30, 31] defined k = 6 as an optimal number of clusters satisfying criteria of algorithm stability, high variance-explained, and reproducibility between independent datasets. The clustered fMRI frames were voxel-wise averaged into CAP maps, and these were then parcellated using the above-mentioned regions by averaging the fMRI activity of the voxels within the region of interest in each CAP map.

### 4.4 Neural network model

Our network model of the mouse cortex is composed of 34 regions (17 for each hemisphere), labelled in Fig. 1A. In turn, each of these regions is composed of one excitatory (E) and one inhibitory (I) population. The excitatory populations are recurrently connected by the 34 × 34 directed anatomical connectivity matrix *J*_*E,E*_. Each entry of *J*_*E,E*_was estimated from the mouse connectome [24] as the number of connections from the entire cortical source region to the unit volume of the cortical target region [50], multiplied by a global scaling coefficient *G*_*E,E*_, which represents the average synaptic efficacy per unit of anatomical connectivity strength and is assumed to be constant across all pairs of regions. Note that, in keeping with previous investigations [23], the matrix *J*_*E,E*_is structured such that the R to R and R to L connections originating from the R hemisphere are respectively identical to the L to L and L to R connections originating from the L hemisphere.

Each excitatory population was connected locally to its corresponding inhibitory population. The weights of the I to E, E to I and I to I connections were collected in the matrices *J*_*E*,*I*_, *J*_*I*,*E*_ and *J*_*I*,*I*_. The values of the entries of these matrices were determined by best fit. The matrices *J*_*I*,*E*_ and *J*_*I*,*I*_ were constructed to have the same value across all regions, whereas the matrix *J*_*E*,*I*_ had values that could be different across regions. This was because in preliminary runs of the model we verified that having region-dependent *J*_*E*,*I*_ seemed to improve the fit quality, whereas having entries of *J*_*I*,*E*_ and *J*_*I*,*I*_ that were region-dependent seemed not to improve fit quality (see also [9]). Since the values of *J*_*E*,*I*_, *J*_*I*,*E*_ and *J*_*I*,*I*_ were determined by best fit rather than by anatomical measures, it was not necessary to include a scaling coefficient representing synaptic efficacy, because this scaling was effectively determined by the best fit.

The mean firing rate of the *i*-th population in the time interval [*t*, *t* + 1) is described by the binary variable *A*_*i*_(*t*), so that *A*_*i*_(*t*) = 0 if the population is silent at time *t*, while *A*_*i*_(*t*) = 1 if it is firing. The spiking activity vector collecting the firing rates of the 68 cortical populations, can switch over time among a set of 2^68^ ≈ 3 × 10^20^ distinct activity patterns. The spiking activity evolves at discrete time instants, where the time step corresponds to the repetition time TR = 1.2*s*. It should be noted that the variations of the model’s spiking activities that are updated from frame to frame should not be interpreted as variations of firing within an integration time constant of the neurons, but rather as time-averaged variations in neural activity on the time scales of the fMRI frame rates.

In each population, the incoming synaptic weights are multiplied by the presynaptic activities at time *t*, to produce the total postsynaptic current. Then, by adding this current to a noise term expressing the net effect of stochastic components of neural activity, we get the mean membrane potential of that population (see Fig. 1B). The membrane potential is passed through a threshold-based activation function, whose output *A*_*i*_(*t* + 1) represents the mean activity of that population at the next time instant. If the membrane potential is below a firing threshold *V*^thr^, then *A*_*i*_(*t* + 1) = 0, otherwise *A*_*i*_(*t* + 1) = 1. Note that the cortical populations do not receive any afferent currents from subcortical regions.

The noise sources in the model are independent and normally distributed, with standard deviation σ, as typically used in whole-brain models (e.g. [5, 10]). These noise sources include all sources that could make neural activity stochastic [80].

To summarize the above with a compact set of equations, we sorted the 68 network nodes so that the excitatory (*E*) populations are labelled by the indexes *i* ∈ {1, ⋯, 34}, while the inhibitory (*I*) populations by the indexes *i* ∈ {35, ⋯, 68}. The mean firing rate of the *i*-th population at time instant *t*, namely *A*_*i*_(*t*) ∈ {0, 1}, is updated at discrete time instants as follows:

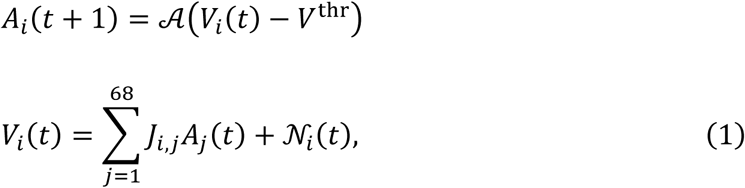

for *i* ∈ {1, ⋯, 68}. The matrix *J*_*i*,*j*_has entries from *J*_*E,E*_, *J*_*E*,*I*_, *J*_*I*,*E*_ or *J*_*I*,*I*_ depending on the index value. In Eq. (1), *V*_*i*_is the mean membrane potential of the *i*-th population, while *A*(⋅) represents the activation function of the network. For our model and for the undirected model discussed in section 2.6, *A*(⋅) is the Heaviside step function:

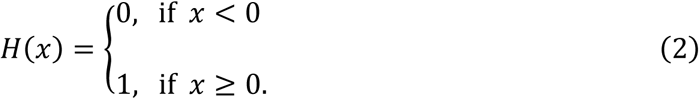

For the linear model reported in section 2.6, *A*(*x*) = *x*. Moreover, ***N***(*t*)^<u>def</u>^ (*N*_1_(*t*), ⋯, *N*_68_(*t*)) is a set of normally distributed sources of neuronal noise, with mean zero and standard deviation σ. The noise terms are spatially and temporally independent, namely Corr (*N*_*i*_(*t*), *N*_*j*_(*s*)) = 0 ∀*i*, *j*, *t*, *s* when *i* ≠ *j* and/or *t* ≠ *s*. Note that, as we focus on cortico-cortical interactions, in Eq. (1) we did not include inputs from subcortical regions.

Based on the finding that fMRI correlates more with Local Field Potentials than with local spiking activity [51, 52], and that LFPs mostly reflect synaptic inputs to pyramidal neurons [51, 53, 54, 81], we modeled the fMRI activity in each cortical region as the total postsynaptic currents on the corresponding excitatory populations. This model provided better fits to the FC values (see section 2.1) than using the spiking activity (not shown).

The model has a set of free parameters, namely *G*_*E,E*_, *J*_*E*,*I*_, *J*_*I*,*E*_, *J*_*I*,*I*_, *V*^thr^, σ. Their values, which are reported in Supplementary Information Table S1, have been obtained by fitting the model to the static values of mean activity and FC of empirical mouse fMRI data, as detailed in section 4.5.

### 4.5 Model fitting procedure

Here we describe how we calculated the values of the free network parameters introduced in section 4.4, namely the global scaling coefficient of the anatomical connectivity (*G*_*E,E*_), the strengths of the I to E, E to I, I to I connections (*J*_*E*,*I*_, *J*_*I*,*E*_, *J*_*I*,*I*_, respectively), the membrane potential firing threshold (*V*^thr^), and the standard deviation of the noise sources (σ).

We derived the best-fit estimates of these parameters by finding the values that minimized the mean squared error between the time-averaged first- and second-order statistics of the empirical mouse fMRI activity and their model predictions. Specifically, the first-order statistic that we considered in the fitting procedure is the relative across-region distribution of the time-averaged fMRI activity, while the second-order statistic is the across-subject averaged functional connectivity matrix. We calculated the first- and second-order statistics over the whole time series, which comprise 450 time points each (9 minutes). Moreover, the empirical statistics have been averaged over 15 mice, while the model statistics over 100 repetitions of the network. In the fitting procedure, we weighed equally in the loss function the first- and second-order terms. We first performed a global coarse-grained grid search and then perfectioned the solution locally in the parameters space with the steepest descent procedure. The values of the best-fit parameters so obtained are listed in Table S1.

### 4.6 Relative across-region distribution of the time-averaged fMRI

The time-averaged mean of the fMRI signals is a first-order statistic that has been typically employed in functional neuroimaging studies to measure the central tendency of the activity [82]. Here, we calculated the relative across-region distribution of the time-averaged fMRI across the 34 cortical regions of the mouse cortex, by subtracting the value of the global signal (that is, the average over all cortical regions of the fMRI signal [9]) from the mean signals of each region and finally by dividing the resulting values by the maximum absolute value across all regions, obtaining a normalized vector whose entries lie in the range [-1, 1].

### 4.7 Static functional connectivity matrix

The static Functional Connectivity (FC) matrix was computed as the across-region Pearson’s correlation estimated over the whole time series and calculated without global signal regression (see Figs. 2D and 2E).

### 4.8 Calculation of the correlation coefficients

We calculated numerically the Pearson’s correlation and its p-value through the function stats.pearsonr() of the Python library Scipy [83].

As is common practice (see e.g. [9, 39]), we calculated the similarity between the full (i.e. 34×34) model and empirical FC matrices, or between symmetric partitions of those matrices (e.g. Figs. 3D, E), by arranging the entries above their main diagonals as vectors, and then by calculating the Pearson’s coefficients to the resulting vectors. For asymmetric partitions (e.g. Figs. 3F, G), we used the whole set of entries to construct the vectors.

In sections 2.3 and S2, we evaluated numerically the similarity between the occupancy probabilities of the basins of attraction by their Spearman’s correlation, through the function stats.spearmanr() of Scipy [83]. The reason for preferring the Spearman’s correlation over other measures of similarity in our analysis of sections 2.3 and S2, is that it is less sensitive to outliers, and the probability distributions in Figs. 4C, F all had one outlier, namely the probability to observe the network state in any oscillatory basin.

### 4.9 Community detection

The community structure of the functional network shown in Fig. 2I was derived using the leading eigenvector method [56], implemented in the function community_leading_eigenvector() of the Python library igraph [84].

### 4.10 Attractors detection

We detected the spiking activity attractors by initializing the network model with a random activity pattern. Then we allowed the pattern to evolve stochastically under the effect of noise for a given number *n* of time steps (typically *n* = 100, so that the network statistics become stationary). At every time step, the spiking activity pattern was updated by solving iteratively Eq. (1) of Methods. After *n* time steps we turned off the noise sources, so that the spiking activity could converge to an attractor of the network without jumping randomly between several attractors. This procedure was then repeated 100,000 times starting from distinct initial patterns, in order to detect the largest number of attractors.

Note that the above algorithm is not guaranteed to find the whole set of attractors, especially when the number of network repetitions is small compared to the total number of attractors. While this algorithm can fully detect the relatively small set of stationary attractors in a few thousands of network repetitions, typically it can reconstruct the prohibitively large set of oscillatory attractors only partially.

### 4.11 Attractor mapping algorithm

In Fig. S1C we sketch the logic of our mapping algorithm. Its main ingredient is a classifier, which labels each time frame of the to-be-analyzed (empirical or model) fMRI time series with the index of the basin of attraction the data point most likely belongs to. The algorithm takes the z-score of the to-be-analyzed 34-dimensional fMRI signal, and compares it (by computing Euclidean distance) with the z-score of the 34-dimensional model fMRI activity, averaged over the basin of each model attractor. This provides a measure of distance between fMRI activity at each time point and each basin of attraction. At every time instant, the classifier labels each time point of the to-be-analyzed fMRI times series with the index of the basin of attraction of the model exhibiting the shortest Euclidean distance (see Fig. S1D). By counting the number of data points across the time series that are assigned to each basin of attraction, we infer the probability of occupation of the basins (blue bars in Figs. 4C, F). To provide a measure of the topography of each estimated basin of attraction that can be compared with the topography of the basins of the network model, the mapping algorithm calculates the mean of the z-score of the to-be-analyzed fMRI signal on all the time points that the classifier assigned to each basin of attraction. Example comparisons are shown in Figs. 4A, D for attractor #17, and in Figs. 4B, E for attractor #13.

### 4.12 Methods to assess non-homotopicity in attractor structure

To assess whether the non-homotopicity in the model attractors had a counterpart in the empirical fMRI data, we used the algorithm of section 4.11 to map the empirical fMRI data either by respecting the differences between non-homotopic attractors (that is, as in Fig. 4F we mapped each fMRI data point into the basins of each of the 32 attractors) or by artificially eliminating the differences between the non-homotopic attractor pairs (this was made by pooling the two basins of attraction of two mirror-symmetric model attractors (e.g. attractors #2, 6) into only one basin prior to the attractor mapping). To compute the effect of enforcing homotopicity on the probability of occupation of the basins of attraction and on the reconstruction of the CAP topography from the model’s attractors, we run the mapping algorithm 100 times, and in each run we simulated both the original model with non-homotopic attractors and the model with enforced homotopic attractor structure 100,000 times. In each of the 100 runs, we evaluated against empirical data the accuracy of the probability of occupation of the basins by using the Spearman’s correlation (as in Fig. 4F), and the accuracy of the CAP reconstructed topography by using the Pearson’s correlation (as in Fig. 5D). Then we performed a Welch’s t-test to assess whether the sets of correlations generated by the two models had equal means.

## References

1. van den Heuvel MP, Hulshoff Pol HE. Exploring the brain network: a review on resting-state fMRI functional connectivity. Eur Neuropsychopharmacol. 2010;20(8):519–34. Epub 2010/05/18. doi: 10.1016/j.euroneuro.2010.03.008. PubMed PMID: 20471808.

2. Rogers BP, Morgan VL, Newton AT, Gore JC. Assessing functional connectivity in the human brain by fMRI. Magn Reson Imaging. 2007;25(10):1347–57. Epub 2007/05/15. doi: 10.1016/j.mri.2007.03.007. PubMed PMID: 17499467; PubMed Central PMCID: PMCPMC2169499.

3. Li K, Guo L, Nie J, Li G, Liu T. Review of methods for functional brain connectivity detection using fMRI. Comput Med Imaging Graph. 2009;33(2):131–9. Epub 2008/12/30. doi: 10.1016/j.compmedimag.2008.10.011. PubMed PMID: 19111443.

4. Deco G, Jirsa VK. Ongoing cortical activity at rest: criticality, multistability, and ghost attractors. J Neurosci. 2012;32(10):3366–75. doi: 10.1523/JNEUROSCI.2523-11.2012. PubMed PMID: 22399758.

5. Deco G, Senden M, Jirsa V. How anatomy shapes dynamics: a semi-analytical study of the brain at rest by a simple spin model. Front Comput Neurosci. 2012;6:68. doi: 10.3389/fncom.2012.00068. PubMed PMID: 23024632.

6. Deco G, Kringelbach ML, Jirsa VK, Ritter P. The dynamics of resting fluctuations in the brain: metastability and its dynamical cortical core. Sci Rep. 2017;7(1):3095. doi: 10.1038/s41598-017-03073-5. PubMed PMID: 28596608.

7. Golos M, Jirsa V, Dauce E. Multistability in Large Scale Models of Brain Activity. PLoS Comput Biol. 2015;11(12):e1004644. doi: 10.1371/journal.pcbi.1004644. PubMed PMID: 26709852.

8. Haimovici A, Tagliazucchi E, Balenzuela P, Chialvo DR. Brain organization into resting state networks emerges at criticality on a model of the human connectome. Phys Rev Lett. 2013;110(17):178101. Epub 2013/05/18. doi: 10.1103/PhysRevLett.110.178101. PubMed PMID: 23679783.

9. Deco G, Ponce-Alvarez A, Hagmann P, Romani GL, Mantini D, Corbetta M. How local excitation-inhibition ratio impacts the whole brain dynamics. J Neurosci. 2014;34(23):7886–98. doi: 10.1523/JNEUROSCI.5068-13.2014. PubMed PMID: 24899711.

10. Deco G, Ponce-Alvarez A, Mantini D, Romani GL, Hagmann P, Corbetta M. Resting-state functional connectivity emerges from structurally and dynamically shaped slow linear fluctuations. J Neurosci. 2013;33(27):11239–52. doi: 10.1523/JNEUROSCI.1091-13.2013. PubMed PMID: 23825427.

11. Ponce-Alvarez A, He BJ, Hagmann P, Deco G. Task-Driven Activity Reduces the Cortical Activity Space of the Brain: Experiment and Whole-Brain Modeling. PLoS Comput Biol. 2015;11(8):e1004445. doi: 10.1371/journal.pcbi.1004445. PubMed PMID: 26317432.

12. Demirtas M, Falcon C, Tucholka A, Gispert JD, Molinuevo JL, Deco G. A whole-brain computational modeling approach to explain the alterations in resting-state functional connectivity during progression of Alzheimer’s disease. Neuroimage Clin. 2017;16:343–54. doi: 10.1016/j.nicl.2017.08.006. PubMed PMID: 28861336.

13. Hansen EC, Battaglia D, Spiegler A, Deco G, Jirsa VK. Functional connectivity dynamics: modeling the switching behavior of the resting state. Neuroimage. 2015;105:525–35. Epub 2014/12/03. doi: 10.1016/j.neuroimage.2014.11.001. PubMed PMID: 25462790.

14. Deco G, Kringelbach ML. Turbulent-like Dynamics in the Human Brain. Cell Rep. 2020;33(10):108471. doi: 10.1016/j.celrep.2020.108471. PubMed PMID: 33296654.

15. Deco G, Sanz Perl Y, Vuust P, Tagliazucchi E, Kennedy H, Kringelbach ML. Rare long-range cortical connections enhance human information processing. Curr Biol. 2021;31(20):4436–48 e5. doi: 10.1016/j.cub.2021.07.064. PubMed PMID: 34437842.

16. Deco G, Van Hartevelt TJ, Fernandes HM, Stevner A, Kringelbach ML. The most relevant human brain regions for functional connectivity: Evidence for a dynamical workspace of binding nodes from whole-brain computational modelling. Neuroimage. 2017;146:197–210. doi: 10.1016/j.neuroimage.2016.10.047. PubMed PMID: 27825955.

17. Glomb K, Ponce-Alvarez A, Gilson M, Ritter P, Deco G. Resting state networks in empirical and simulated dynamic functional connectivity. Neuroimage. 2017;159:388–402. doi: 10.1016/j.neuroimage.2017.07.065. PubMed PMID: 28782678.

18. Rabuffo G, Fousek J, Bernard C, Jirsa V. Neuronal Cascades Shape Whole-Brain Functional Dynamics at Rest. eNeuro. 2021;8(5):e0283–21.2021. doi: 10.1523/ENEURO.0283-21.2021. PubMed PMID: 34583933.

19. Arbabyazd L, Shen K, Wang Z, Hofmann-Apitius M, Ritter P, McIntosh AR, et al. Virtual Connectomic Datasets in Alzheimer’s Disease and Aging Using Whole-Brain Network Dynamics Modelling. eneuro. 2021;8(4):ENEURO.0475-20.2021. doi: 10.1523/ENEURO.0475-20.2021.

20. Zerbi V, Pagani M, Markicevic M, Matteoli M, Pozzi D, Fagiolini M, et al. Brain mapping across 16 autism mouse models reveals a spectrum of functional connectivity subtypes. Mol Psychiatry. 2021;26:7610–20. doi: 10.1038/s41380-021-01245-4. PubMed PMID: 34381171.

21. Pagani M, Barsotti N, Bertero A, Trakoshis S, Ulysse L, Locarno A, et al. mTOR-related synaptic pathology causes autism spectrum disorder-associated functional hyperconnectivity. Nature Communications. 2021;12(1):6084. doi: 10.1038/s41467-021-26131-z. PubMed PMID: 34667149.

22. Rocchi F, Canella C, Noei S, Gutierrez-Barragan D, Coletta L, Galbusera A, et al. Increased fMRI connectivity upon chemogenetic inhibition of the mouse prefrontal cortex. Nature Communications. 2022;13:1056. doi: doi.org/10.1038/s41467-022-28591-3

23. Coletta L, Pagani M, Whitesell JD, Harris JA, Bernhardt B, Gozzi A. Network structure of the mouse brain connectome with voxel resolution. Sci Adv. 2020;6(51):eabb7187. doi: 10.1126/sciadv.abb7187. PubMed PMID: 33355124.

24. Knox JE, Harris KD, Graddis N, Whitesell JD, Zeng H, Harris JA, et al. High-resolution data-driven model of the mouse connectome. Netw Neurosci. 2019;3(1):217–36. Epub 20181201. doi: 10.1162/netn_a_00066. PubMed PMID: 30793081.

25. Oh SW, Harris JA, Ng L, Winslow B, Cain N, Mihalas S, et al. A mesoscale connectome of the mouse brain. Nature. 2014;508(7495):207-14. doi: 10.1038/nature13186.

26. Bullmore E, Sporns O. Complex brain networks: graph theoretical analysis of structural and functional systems. Nature Reviews Neuroscience. 2009;10(3):186–98. doi: 10.1038/nrn2575.

27. Straathof M, Sinke MR, Dijkhuizen RM, Otte WM. A systematic review on the quantitative relationship between structural and functional network connectivity strength in mammalian brains. J Cereb Blood Flow Metab. 2019;39(2):189–209. doi: 10.1177/0271678X18809547. PubMed PMID: 30375267.

28. Grandjean J, Zerbi V, Balsters JH, Wenderoth N, Rudin M. Structural Basis of Large-Scale Functional Connectivity in the Mouse. J Neurosci. 2017;37(34):8092–101. doi: 10.1523/JNEUROSCI.0438-17.2017. PubMed PMID: 28716961.

29. Stafford JM, Jarrett BR, Miranda-Dominguez O, Mills BD, Cain N, Mihalas S, et al. Large-scale topology and the default mode network in the mouse connectome. Proc Natl Acad Sci USA. 2014;111(52):18745–50. doi: 10.1073/pnas.1404346111.

30. Gutierrez-Barragan D, Basson MA, Panzeri S, Gozzi A. Infraslow State Fluctuations Govern Spontaneous fMRI Network Dynamics. Current Biology. 2019;29(14):2295–306 e5. doi: 10.1016/j.cub.2019.06.017. PubMed PMID: 31303490.

31. Gutierrez-Barragan D, Singh NA, Alvino FG, Coletta L, Rocchi F, De Guzman E, et al. Unique spatiotemporal fMRI dynamics in the awake mouse brain. Current Biology. 2022;32:631–44. doi: 10.1016/j.cub.2021.12.015.

32. Liska A, Galbusera A, Schwarz AJ, Gozzi A. Functional connectivity hubs of the mouse brain. Neuroimage. 2015;115:281–91. doi: 10.1016/j.neuroimage.2015.04.033. PubMed PMID: 25913701.

33. Grandjean J, Canella C, Anckaerts C, Ayranci G, Bougacha S, Bienert T, et al. Common functional networks in the mouse brain revealed by multi-centre resting-state fMRI analysis. Neuroimage. 2020;205:116278. doi: 10.1016/j.neuroimage.2019.116278. PubMed PMID: 31614221.

34. Liu X, Duyn JH. Time-varying functional network information extracted from brief instances of spontaneous brain activity. Proc Natl Acad Sci U S A. 2013;110(11):4392–7. doi: 10.1073/pnas.1216856110. PubMed PMID: 23440216.

35. Liu X, Zhang N, Chang C, Duyn JH. Co-activation patterns in resting-state fMRI signals. Neuroimage. 2018;180(Pt B):485–94. doi: 10.1016/j.neuroimage.2018.01.041. PubMed PMID: 29355767.

36. Huang Z, Zhang J, Wu J, Mashour GA, Hudetz AG. Temporal circuit of macroscale dynamic brain activity supports human consciousness. Sci Adv. 2020;6(11). Epub 2020/03/21. doi: 10.1126/sciadv.aaz0087. PubMed PMID: 32195349.

37. Gutierrez-Barragan D, Ramirez JSB, Panzeri S, Xu T, Gozzi A. Evolutionarily conserved fMRI network dynamics in the mouse, macaque, and human brain. Nature Communications. 2024;15(1):8518. doi: 10.1038/s41467-024-52721-8.

38. Melozzi F, Woodman MM, Jirsa VK, Bernard C. The Virtual Mouse Brain: A Computational Neuroinformatics Platform to Study Whole Mouse Brain Dynamics. eNeuro. 2017;4(3):e0111–17.2017. Epub 20170628. doi: 10.1523/ENEURO.0111-17.2017. PubMed PMID: 28664183; PubMed Central PMCID: PMCPMC5489253.

39. Melozzi F, Bergmann E, Harris JA, Kahn I, Jirsa V, Bernard C. Individual structural features constrain the mouse functional connectome. Proc Natl Acad Sci USA. 2019;116:26961–9. Epub 20191211. doi: 10.1073/pnas.1906694116. PubMed PMID: 31826956; PubMed Central PMCID: PMCPMC6936369.

40. Siu PH, Müller E, Zerbi V, Aquino K, Fulcher BD. Extracting Dynamical Understanding From Neural-Mass Models of Mouse Cortex. Frontiers in Computational Neuroscience. 2022;16:847336. doi: 10.3389/fncom.2022.847336.

41. Raut RV, Snyder AZ, Mitra A, Yellin D, Fujii N, Malach R, et al. Global waves synchronize the brain’s functional systems with fluctuating arousal. Science Advances. 7(30):eabf2709. doi: 10.1126/sciadv.abf2709.

42. Zamani Esfahlani F, Jo Y, Faskowitz J, Byrge L, Kennedy DP, Sporns O, et al. High-amplitude cofluctuations in cortical activity drive functional connectivity. Proc Natl Acad Sci USA. 2020;117(45):28393–401. doi: 10.1073/pnas.2005531117. PubMed PMID: 33093200.

43. Munn BR, Muller EJ, Wainstein G, Shine JM. The ascending arousal system shapes neural dynamics to mediate awareness of cognitive states. Nature Communications. 2021;12(1):6016. Epub 2021/10/16. doi: 10.1038/s41467-021-26268-x. PubMed PMID: 34650039; PubMed Central PMCID: PMCPMC8516926.

44. Shine JM. Neuromodulatory Influences on Integration and Segregation in the Brain. Trends in Cognitive Sciences. 2019;23(7):572–83. doi: doi.org/10.1016/j.tics.2019.04.002.

45. Whitesell JD, Liska A, Coletta L, Hirokawa KE, Bohn P, Williford A, et al. Regional, Layer, and Cell-Type-Specific Connectivity of the Mouse Default Mode Network. Neuron. 2021;109(3):545–59 e8. doi: 10.1016/j.neuron.2020.11.011. PubMed PMID: 33290731.

46. Fasoli D, Panzeri S. Optimized brute-force algorithms for the bifurcation analysis of a binary neural network model. Phys Rev E. 2019;99(1-1):012316. doi: 10.1103/PhysRevE.99.012316. PubMed PMID: 30780305.

47. Amari SI. Learning Patterns and Pattern Sequences by Self-Organizing Nets of Threshold Elements. IEEE Transactions on Computers. 1972;C-21(11):1197-206. doi: 10.1109/T-C.1972.223477.

48. Little WA. The existence of persistent states in the brain. Mathematical Biosciences. 1974;19(1):101–20. doi: 10.1016/0025-5564(74)90031-5.

49. Peretto P. Collective properties of neural networks: A statistical physics approach. Biological Cybernetics. 1984;50(1):51–62. doi: 10.1007/BF00317939.

50. Rubinov M, Ypma RJ, Watson C, Bullmore ET. Wiring cost and topological participation of the mouse brain connectome. Proc Natl Acad Sci USA. 2015;112(32):10032–7. doi: 10.1073/pnas.1420315112. PubMed PMID: 26216962.

51. Logothetis NK. What we can do and what we cannot do with fMRI. Nature. 2008;453(7197):869-78. Epub 2008/06/13. doi: 10.1038/nature06976. PubMed PMID: 18548064.

52. Logothetis NK, Pauls J, Augath M, Trinath T, Oeltermann A. Neurophysiological investigation of the basis of the fMRI signal. Nature. 2001;412(6843):150-7. Epub 2001/07/13. doi: 10.1038/35084005. PubMed PMID: 11449264.

53. Einevoll GT, Kayser C, Logothetis NK, Panzeri S. Modelling and analysis of local field potentials for studying the function of cortical circuits. Nat Rev Neurosci. 2013;14(11):770–85. doi: 10.1038/nrn3599. PubMed PMID: 24135696.

54. Mazzoni A, Panzeri S, Logothetis NK, Brunel N. Encoding of naturalistic stimuli by local field potential spectra in networks of excitatory and inhibitory neurons. PLoS Comput Biol. 2008;4(12):e1000239. doi: 10.1371/journal.pcbi.1000239. PubMed PMID: 19079571.

55. Sforazzini F, Schwarz AJ, Galbusera A, Bifone A, Gozzi A. Distributed BOLD and CBV-weighted resting-state networks in the mouse brain. Neuroimage. 2014;87:403–15. Epub 2013/10/02. doi: 10.1016/j.neuroimage.2013.09.050. PubMed PMID: 24080504.

56. Newman MEJ. Finding community structure in networks using the eigenvectors of matrices. Physical Review E. 2006;74(3):036104. doi: 10.1103/PhysRevE.74.036104.

57. Fox MD, Snyder AZ, Vincent JL, Corbetta M, Van Essen DC, Raichle ME. The human brain is intrinsically organized into dynamic, anticorrelated functional networks. Proc Natl Acad Sci U S A. 2005;102(27):9673–8. doi: 10.1073/pnas.0504136102. PubMed PMID: 15976020.

58. Fox MD, Zhang D, Snyder AZ, Raichle ME. The Global Signal and Observed Anticorrelated Resting State Brain Networks. Journal of Neurophysiology. 2009;101(6):3270–83. doi: 10.1152/jn.90777.2008.

59. Murphy K, Birn RM, Handwerker DA, Jones TB, Bandettini PA. The impact of global signal regression on resting state correlations: are anti-correlated networks introduced? Neuroimage. 2009;44(3):893–905. Epub 20081011. doi: 10.1016/j.neuroimage.2008.09.036. PubMed PMID: 18976716; PubMed Central PMCID: PMCPMC2750906.

60. Weissenbacher A, Kasess C, Gerstl F, Lanzenberger R, Moser E, Windischberger C. Correlations and anticorrelations in resting-state functional connectivity MRI: a quantitative comparison of preprocessing strategies. Neuroimage. 2009;47(4):1408–16. doi: 10.1016/j.neuroimage.2009.05.005. PubMed PMID: 19442749.

61. Lurie DJ, Kessler D, Bassett DS, Betzel RF, Breakspear M, Kheilholz S, et al. Questions and controversies in the study of time-varying functional connectivity in resting fMRI. Network Neuroscience. 2020;4(1):30–69. doi: 10.1162/netn_a_00116.

62. Yang H, Zhang H, Di X, Wang S, Meng C, Tian L, et al. Reproducible coactivation patterns of functional brain networks reveal the aberrant dynamic state transition in schizophrenia. NeuroImage. 2021;237:118193.

63. Hopfield JJ. Neural networks and physical systems with emergent collective computational abilities. Proc Natl Acad Sci U S A. 1982;79(8):2554–8. doi: 10.1073/pnas.79.8.2554. PubMed PMID: 6953413.

64. Strocchi F. Spontaneous Symmetry Breaking. Lecture Notes in Physics. 2008;732:9–11. doi: 10.1007/978-3-540-73593-9_2.

65. Bernstein J. Spontaneous symmetry breaking, gauge theories, the Higgs mechanism and all that. Reviews of Modern Physics. 1974;46(1):7–48. doi: 10.1103/RevModPhys.46.7.

66. Fasoli D, Cattani A, Panzeri S. The Complexity of Dynamics in Small Neural Circuits. PLoS Comput Biol. 2016;12(8):e1004992. doi: 10.1371/journal.pcbi.1004992. PubMed PMID: 27494737.

67. Betzel RF, Bassett DS. Specificity and robustness of long-distance connections in weighted, interareal connectomes. Proc Natl Acad Sci USA. 2018;115(21):E4880–E9. doi: 10.1073/pnas.1720186115.

68. Khanjanianpak M, Azimi-Tafreshi N, Valizadeh A. Emergence of complex oscillatory dynamics in the neuronal networks with long activity time of inhibitory synapses. iScience. 2024;27(4). doi: 10.1016/j.isci.2024.109401.

69. Kringelbach ML, Cruzat J, Cabral J, Knudsen GM, Carhart-Harris R, Whybrow PC, et al. Dynamic coupling of whole-brain neuronal and neurotransmitter systems. Proc Natl Acad Sci U S A. 2020;117(17):9566–76. doi: 10.1073/pnas.1921475117. PubMed PMID: 32284420.

70. Deco G, Cabral J, Saenger VM, Boly M, Tagliazucchi E, Laufs H, et al. Perturbation of whole-brain dynamics in silico reveals mechanistic differences between brain states. NeuroImage. 2018;169:46–56. doi: 10.1016/j.neuroimage.2017.12.009.

71. Jobst BM, Hindriks R, Laufs H, Tagliazucchi E, Hahn G, Ponce-Alvarez A, et al. Increased Stability and Breakdown of Brain Effective Connectivity During Slow-Wave Sleep: Mechanistic Insights from Whole-Brain Computational Modelling. Scientific Reports. 2017;7(1):4634. doi: 10.1038/s41598-017-04522-x.

72. Ding X, Froudist-Walsh S, Jaramillo J, Jiang J, Wang X-J. Cell type-specific connectome predicts distributed working memory activity in the mouse brain. eLife. 2024;13:e85442. doi: 10.7554/eLife.85442.

73. Van de Ville D. Brain Dynamics: Global Pulse and Brain State Switching. Current Biology. 2019;29(14):R690-R2. doi: 10.1016/j.cub.2019.06.006.

74. Power JD, Mitra A, Laumann TO, Snyder AZ, Schlaggar BL, Petersen SE. Methods to detect, characterize, and remove motion artifact in resting state fMRI. Neuroimage. 2014;84:320–41. Epub 2013/09/03. doi: 10.1016/j.neuroimage.2013.08.048. PubMed PMID: 23994314; PubMed Central PMCID: PMCPMC3849338.

75. Chang C, Leopold DA, Scholvinck ML, Mandelkow H, Picchioni D, Liu X, et al. Tracking brain arousal fluctuations with fMRI. Proc Natl Acad Sci USA. 2016;113(16):4518–23. doi: 10.1073/pnas.1520613113. PubMed PMID: 27051064.

76. Yang GJ, Murray JD, Repovs G, Cole MW, Savic A, Glasser MF, et al. Altered global brain signal in schizophrenia. Proc Natl Acad Sci USA. 2014;111(20):7438–43. Epub 2014/05/07. doi: 10.1073/pnas.1405289111. PubMed PMID: 24799682.

77. Liu X, de Zwart JA, Scholvinck ML, Chang C, Ye FQ, Leopold DA, et al. Subcortical evidence for a contribution of arousal to fMRI studies of brain activity. Nature Communications. 2018;9(1):395. doi: 10.1038/s41467-017-02815-3. PubMed PMID: 29374172.

78. Wang X-J. Probabilistic Decision Making by Slow Reverberation in Cortical Circuits. Neuron. 2002;36(5):955–68. doi: 10.1016/S0896-6273(02)01092-9.

79. Mejías JF, Wang X-J. Mechanisms of distributed working memory in a large-scale network of macaque neocortex. eLife. 2022;11:e72136. doi: 10.7554/eLife.72136.

80. Faisal AA, Selen LPJ, Wolpert DM. Noise in the nervous system. Nature Reviews Neuroscience. 2008;9(4):292–303. doi: 10.1038/nrn2258.

81. Martínez-Cañada P, Ness TV, Einevoll GT, Fellin T, Panzeri S. Computation of the electroencephalogram (EEG) from network models of point neurons. PLOS Computational Biology. 2021;17(4):e1008893. doi: 10.1371/journal.pcbi.1008893.

82. Grady CL, Garrett DD. Understanding variability in the BOLD signal and why it matters for aging. Brain Imaging Behav. 2014;8(2):274–83. doi: 10.1007/s11682-013-9253-0. PubMed PMID: 24008589.

83. Virtanen P, Gommers R, Oliphant TE, Haberland M, Reddy T, Cournapeau D, et al. SciPy 1.0: fundamental algorithms for scientific computing in Python. Nature Methods. 2020;17(3):261–72. doi: 10.1038/s41592-019-0686-2. PubMed PMID: 32015543.

84. Csardi G, Nepusz T. The Igraph Software Package for Complex Network Research. International Journal of Complex Systems. 2005;23:1695.

