## Supplemental Information for "Attractor dynamics of a whole-cortex network model predicts emergence and structure of fMRI co-activation patterns in the mouse brain"

### Supplementary Results

#### S1 Two-population toy model of formation of non-homotopic attractors through spontaneous symmetry breaking

To explain the formation of the non-homotopicity of attractors in the presence of inter-hemispheric excitatory coupling, we study a minimal bi-hemispheric model with two anatomically identical regions in the two hemispheres, which are recursively connected by intra and inter-hemispheric anatomical connections. Due to the left-right symmetry of the model, the network always exhibits homotopic solutions. However, when the intra- and inter-hemispheric couplings are in a specific range of values, the network has also two non-homotopic attractors. Interestingly, this phenomenon occurs also when the two hemispheres are anatomically connected by relatively strong inter-hemispheric connections. Therefore, contrary to intuition, the presence of inter-hemispheric connections with large synaptic weight allows the two hemispheres to effectively exchange information, but it does not necessarily force them to synchronize. This is the hallmark of the non-linear phenomenon known as Spontaneous Symmetry Breaking (SSB).

In what follows, we perform an analytical study of SSB in our bi-hemispheric toy model. The model is composed of two identical populations with binary firing rates and firing threshold $V^{\mathrm{thr}}$, representing the excitatory populations in two homologous regions of the mouse cortex, e.g. lVIS and rVIS. Each population projects two kinds of anatomical connections: the first has weight $J_{\mathrm{intra}} > 0$ and links the population to itself, while the second has weight $J_{\mathrm{inter}} > 0$ and connects the population to the other one in the model. In other words, $J_{\mathrm{intra}}$ and $J_{\mathrm{inter}}$ are free parameters that describe the strength of the intra and inter-hemispheric connections, respectively. In what follows, we prove that SSB occurs in this bi-hemispheric network whenever $J_{\mathrm{intra}} \geq V^{\mathrm{thr}}$ and $J_{\mathrm{inter}} < V^{\mathrm{thr}}$. Since we want to investigate how SSB leads to the inter-hemispheric non-homotopicity of the neural attractors, regardless of the differences in the neural activity that can arise in the two hemispheres because of the independent sources of noise, in our investigation we set to zero the amplitude of the noise fluctuations. For this reason, the model satisfies the following set of equations (compare with Eq. (1) in the main text for $\sigma = 0$ and $\mathcal{A}(\cdot) = H(\cdot)$):

$$\begin{cases} \mathrm{V}_L(t+1) = J_{\mathrm{intra}} H\left(\mathrm{V}_L(t) - V^{\mathrm{thr}}\right) + J_{\mathrm{inter}} H\left(\mathrm{V}_R(t) - V^{\mathrm{thr}}\right) \\ \mathrm{V}_R(t+1) = J_{\mathrm{intra}} H\left(\mathrm{V}_R(t) - V^{\mathrm{thr}}\right) + J_{\mathrm{inter}} H\left(\mathrm{V}_L(t) - V^{\mathrm{thr}}\right), \end{cases} \tag{S1}$$

where $\mathrm{V}_L$ and $\mathrm{V}_R$ are the model membrane potentials of the cortical regions in the left and right hemispheres. Note that Eq. (S1) is inter-hemispherically homotopic, namely it is invariant under the exchange of $\mathrm{V}_L$ and $\mathrm{V}_R$.

For simplicity, in our investigation we focus on the stationary solutions of Eq. (S1), since they suffice for our purposes. By imposing the stationarity condition $\mathrm{V}_x(t+1) = \mathrm{V}_x(t)$, for $x \in \{L, R\}$, into Eq. (S1), whenever $V^{\mathrm{thr}} > 0$ we get:

- $\forall J_{\mathrm{intra}}, J_{\mathrm{inter}}$, $(\mathrm{V}_L, \mathrm{V}_R) = (0, 0)$ is a homotopic solution to Eq. (S1) (note that in this case both the regions are silent, i.e. $H(\mathrm{V}_L - V^{\mathrm{thr}}) = H(\mathrm{V}_R - V^{\mathrm{thr}}) = 0$);
- if $J_{\mathrm{intra}} + J_{\mathrm{inter}} \geq V^{\mathrm{thr}}$, $(\mathrm{V}_L, \mathrm{V}_R) = (J_{\mathrm{intra}} + J_{\mathrm{inter}}, J_{\mathrm{intra}} + J_{\mathrm{inter}})$ is a homotopic solution to Eq. (S1) (in this case both the regions are firing);
- if $J_{\mathrm{intra}} \geq V^{\mathrm{thr}}$ and $J_{\mathrm{inter}} < V^{\mathrm{thr}}$, $(\mathrm{V}_L, \mathrm{V}_R) = (J_{\mathrm{intra}}, J_{\mathrm{inter}})$ and $(\mathrm{V}_L, \mathrm{V}_R) = (J_{\mathrm{inter}}, J_{\mathrm{intra}})$ are non-homotopic solutions to Eq. (S1) (one region is firing, while the other is silent).

These results show that, for specific values of the network parameters and regardless of the inter-hemispheric homotopicity of Eq. (S1), the model admits a set of non-homotopic solutions, which are the hallmark of SSB. Specifically, when we focus on the main diagonals of the intra- and inter-hemispheric portions of the matrix $J_{E,E}$ in the mouse connectome (see Fig. 1A), which, similarly to $J_{\mathrm{intra}}$ and $J_{\mathrm{inter}}$, represent the strength of intra-regional and homotopic (i.e. between homologous regions across the two hemispheres) connections respectively, the conditions for the emergence of SSB are satisfied in MO, SS, AUD, VIS and RSP. Therefore, according to our study, these regions are expected to drive the formation of inter-hemispheric non-homotopicity in the mouse brain through SSB. Interestingly, they can also induce non-homotopicity in other cortical regions through the inter-regional and non-homotopic connections (i.e. the off-diagonal entries of the intra- and inter-hemispheric portions of the matrix $J_{E,E}$), therefore leading to the formation of the complex non-homotopic attractors that we listed in Table S2 and Fig. 5B.

Generally, when we turn on the noise sources (i.e. for $\sigma > 0$) and whenever $J_{\mathrm{intra}} \geq V^{\mathrm{thr}}$ and $J_{\mathrm{inter}} < V^{\mathrm{thr}}$, the membrane potentials $\mathrm{V}_L$ and $\mathrm{V}_R$ affect each other through the inter-hemispheric connections, while fluctuating stochastically between the homotopic attractors and the two spatially opposite non-homotopic attractors that we described above. For this reason, SSB is observed only over timescales of the order of few frames, and therefore of seconds. Inter-hemispheric non-homotopicity is obscured when the network statistics are calculated over longer timescales, because in this case the network spends approximately the same amount of time in the two non-homotopic attractors, so that the frame-by-frame differences in the activity of the two hemispheres are averaged away.

**S2 Further studies of the correspondence between attractor dynamics predicted by the model and the structure of fMRI dynamics**

To validate our mapping algorithm, we embedded it with the attractor structure of the best-fit model. Then, we asked the algorithm to map on those attractors the time series generated from the network model run with parameters different from the best-fit ones. We modified parametrically the connectivity value of the model by modulating the degree of inter-hemispheric connectivity by a global scaling coefficient $W$. When varying the connectivity, the model still gave rise to attractor dynamics, but the number of attractors and their topography varied with the degree of inter-hemispheric connectivity. When the algorithm allocated the fMRI series of the non-best-fit models to the set of attractors of the model with the best-fit inter-hemispheric connectivity, the reconstructed distribution showed a lower match with that of the embedded attractors compared to the time series generated by the model with the best-fit coupling $W = 1$ (Fig. S2A).

We obtained similar results when we increased the threshold $T$ to produce sparser versions of the connectome (Fig. S2B), and when we modified the local coupling strength between the excitatory (E) and inhibitory (I) populations of the model by a global scaling coefficient $z$ (Fig. S2C). As expected, the mapping algorithm applied on the fMRI time series simulated from our network model with the non-best-fit sparsity or E-I coupling, did not reconstruct the distribution of the embedded best-fit attractors as well as when it is applied on the time series generated by the model with best-fit parameters $T = 0$ and $z = 1$.

Next, to support the notion that the empirical fMRI data are compatible with attractor dynamics features that are specific to those of the best-fit model, we asked the mapping algorithm to allocate the empirical fMRI signals to the different set of attractors predicted by the model with the non-best-fit inter-hemispheric connectivity, sparsity of connections, or E-I coupling. In other words, for each value of $W$, $T$ and $z$, we embedded the mapping algorithm with the specific set of spiking activity attractors generated by the model with the new set of parameters. The probability distribution of the basins reconstructed by the algorithm did not match the new distribution of the basins of spiking activity as well as the case when the empirical fMRI data are mapped on the model with the best-fit parameters (Figs. S2D-F).

Thus, the empirical fMRI time series are more compatible with attractor dynamics of the specific form generated by the best-fit model. It is important to observe that this result was obtained despite the model's parameters were not fitted to maximize the similarity between the attractor structure predicted by the model and the results generated by the mapping algorithm.

We finally asked whether attractor dynamics features that are specific to those of the best-fit model could explain the emergence of CAPs in empirical fMRI data better than the features of the non-best-fit-model attractors. The match between the topography of empirical and model CAPs was maximal when using as model parameters $W$, $T$ and $z$ those determined by best fit, and computing this match with a different set of parameter values produced a weaker match (Figs. S2G-I).

**S3 Mechanistic study of the effect of weak vs strong anatomical connections**

In section 2.4 we showed that the removal of the strongest anatomical connections in the mouse connectome, obtained by gradually increasing the anatomical threshold $T$, leads to a considerable loss of stationary attractors in our network model, while the removal of either weak or strong connections significantly reduces the number of oscillatory attractors (see Fig. 6B). In what follows, we explain mechanistically this phenomenon.

Since the spiking activity attractors are an intrinsic network property that does not depend on noise, in this section we will compute attractor properties by setting $\sigma = 0$ in Eq. (1). Then, we define the sequence of activity patterns $\mathbb{A}(0,\mathcal{T}) \stackrel{\text{def}}{=} \boldsymbol{A}^{(0)} \to \boldsymbol{A}^{(1)} \to \cdots \to \boldsymbol{A}^{(\mathcal{T})}$, for some $\mathcal{T}$ such that $1 \le \mathcal{T} \le 2^{68}$, and we introduce the set $\Gamma_x^{(j)} \stackrel{\text{def}}{=} \left\{i \in \{1,\cdots,68\}\text{: } A_i^{(j)} = x\right\}$, for $x \in \{0,1\}$. By adapting the results of Ref. [46] to our network model, we get that the sequence $\mathbb{A}(0,\mathcal{T})$ is a solution of Eq. (1) in the time range $[0,\mathcal{T}]$ (namely $\boldsymbol{A}(t = j) = \boldsymbol{A}^{(j)}$ for $t = 0,\cdots,\mathcal{T}$), if and only if:

$$L_{\mathbb{A}} \stackrel{\text{def}}{=} \max_{j\in\mathfrak{T}_1}\left(\max_{i\in\Gamma_1^{(j+1)}} \mathfrak{I}_i^{(j)}\right) \le 0, \quad U_{\mathbb{A}} \stackrel{\text{def}}{=} \min_{j\in\mathfrak{T}_0}\left(\min_{i\in\Gamma_0^{(j+1)}} \mathfrak{I}_i^{(j)}\right) > 0, \tag{S2}$$

where:

$$\mathfrak{T}_x \overset{\text{def}}{=} \left\{ j \in \mathfrak{T} : \ \Gamma_x^{(j+1)} \neq \emptyset \right\}, \qquad \mathfrak{T} \overset{\text{def}}{=} \{0, \cdots, \mathcal{T}-1\}$$

$$\mathfrak{I}_i^{(j)} \overset{\text{def}}{=} V^{\text{thr}} - \sum_{k=1}^{68} J_{i,k} A_k^{(j)}.$$

When Eq. (S2) is not satisfied, $\mathbb{A}(0,\mathcal{T})$ is not a solution to Eq. (1), and therefore it turns into a different sequence of activity patterns in the time range $[0,\mathcal{T}]$.

In the specific case when $\boldsymbol{A}^{(0)} = \boldsymbol{A}^{(\mathcal{T})}$, the sequence $\mathbb{A}(0,\mathcal{T})$ represents an oscillatory attractor with period $\mathcal{T}$. Moreover, whenever $\mathcal{T} = 1$, the periodic sequence corresponds to a stationary attractor, since in this case the spiking activity in each cortical region stays constant over time.

In this section, we study how the stability of the attractors changes when we gradually remove anatomical connections from the network model, by increasing the anatomical threshold $T$. Our analysis revealed that the $U_{\mathbb{A}}$ of each network attractor is weakly affected by $T$ (result not shown), therefore in what follows we focus on $L_{\mathbb{A}}$ only.

When we gradually increase the anatomical threshold $T$, several entries $J_{i,k}$ are set to zero, therefore the term $\mathfrak{I}_i^{(j)}$ in Eq. (S2) becomes larger and larger, and the likeliness to get $L_{\mathbb{A}} > 0$ (which would cause the annihilation of the attractor $\mathbb{A}$) increases accordingly. Because of the function $\max_{j\in\mathfrak{T}_1}(\cdot)$ in the formula of $L_{\mathbb{A}}$ (see Eq. (S2)), this likeliness is larger for oscillations with large period $\mathcal{T}$. This implies that the stationary states ($\mathcal{T} = 1$) are weakly affected by the removal of the weakest anatomical connections, so that they annihilate only after removing strong anatomical connections (blue curve in Fig. 6B).

On the other hand, the oscillatory attractors in our network model have period $\mathcal{T} = 4$, therefore even the removal of the weakest anatomical connections can destabilize them (red curve in Fig. 6B).

The blue (respectively, red) bars in Fig. S4 illustrate how the distribution of values of $L_{\mathbb{A}}$ across the stationary (respectively, oscillatory) attractors of our model with the best-fit parameters changes when we increase the anatomical threshold gradually from $T = 0$ (Fig. S4A) to, e.g., $T = 0.03$ (Fig. S4B) and $T = 0.07$ (Fig. S4C). Note that it is always $L_{\mathbb{A}} < 0$ for all the blue bars, therefore ensuring the stability of the 31 stationary attractors of the model (compare with the blue curve in Fig. 6B, which exhibits a constant number of stationary attractors up to $T = 0.07$). On the other hand, for $T = 0.03$ and $T = 0.07$, some of the red bars have $L_{\mathbb{A}} > 0$ and therefore become unstable. This result confirms that, unlike the stationary attractors, the oscillatory attractors are affected also by the removal of the weakest anatomical connections.

#### S4 Mechanistic study of the effect of local excitation-inhibition coupling on the stationary attractors of the model

In this section, we explain the neural mechanism underlying the decrease in the number of stationary attractors that we observed by increasing the coupling coefficient $z$ between the excitatory and inhibitory populations of the model (blue curve in Fig. 6C). Similarly to sections S1 and S3, we set to zero the amplitude of the noise fluctuations to perform this investigation.

For a chosen value of the coefficient $z$, the stationary attractors of our model have the form $\boldsymbol{A}^{(p,z)} = \left[\boldsymbol{A}_E^{(p,z)}, \boldsymbol{A}_I^{(p,z)}\right]$ for $p \in \left\{1, \cdots, M^{(z)}\right\}$, where $M^{(z)}$ is the total number of stationary attractors, while $\boldsymbol{A}_E^{(p,z)}$ and $\boldsymbol{A}_I^{(p,z)}$ are the 34-dimensional binary activity patterns of the excitatory and the inhibitory populations in the network, respectively. According to Eq. (1) with $\sigma = 0$ and $\mathcal{A}(\cdot) = H(\cdot)$, the entries $A_{E,i}^{(p,z)}$ and $A_{I,i}^{(p,z)}$ of the two vectors satisfy the following set of equations:

$$\begin{cases} A_{E,i}^{(p,z)} = H\left(\sum_{j=1}^{34} \left[J_{E,E}\right]_{i,j} A_{E,j}^{(p,z)} + z\left[J_{E,I}\right]_i A_{I,i}^{(p,z)} - V^{\text{thr}}\right) \\ \\ A_{I,i}^{(p,z)} = H\left(z\left[J_{I,E}\right]_i A_{E,i}^{(p,z)} + \left[J_{I,I}\right]_i A_{I,i}^{(p,z)} - V^{\text{thr}}\right), \end{cases} \tag{S3}$$

$\forall i \in \{1, \cdots, 34\}$. Given the values of the best-fit parameters in Table S1, for $z \geq \frac{V^{\text{thr}} - [J_{I,I}]_i}{[J_{I,E}]_i} = 0.38$ the second identity in Eq. (S3) has solution $A_{I,i}^{(p,z)} = A_{E,i}^{(p,z)}$. Therefore, for $z \geq 0.38$, the first identity can be rewritten as follows:

$$A_{E,i}^{(p,z)} = H\left(\sum_{j=1}^{34} \left[J_{E,E}\right]_{i,j} A_{E,j}^{(p,z)} + z\left[J_{E,I}\right]_i A_{E,i}^{(p,z)} - V^{\text{thr}}\right), \qquad \forall i \in \{1, \cdots, 34\}. \tag{S4}$$

In what follows, we study the variation in the number of stationary attractors specifically in this range of the scaling coefficient $z$ (note that the case $0 < z < 0.38$, being very different from the best-fit value $z = 1$, is not considered here).

To describe the variation in the number of stationary attractors as function of $z$, it is useful to study how the number of solutions to Eq. (S4) changes with respect to a reference network, when we increase $z$. For our purpose, it is convenient to choose the case $z = 0$ (excitatory populations fully disconnected from their local inhibitory populations) as reference point, and to study the number of solutions to Eq. (S4) when $z$ increases above 0.38.

Let us assume that $A_{E,i}^{(p,0)}$, for $p \in \left\{1, \cdots, M^{(0)}\right\}$, are the solutions to Eq. (S4) for $z = 0$. A necessary condition for $A_{E,i}^{(p,0)}$ to be a solution to Eq. (S4) also for some $z \geq 0.38$, is:

$$z \leq z^{(p)} \stackrel{\text{def}}{=} \min_{i \in \Gamma^{(p)}} \left(\frac{V^{\text{thr}} - \sum_{j=1}^{34} \left[J_{E,E}\right]_{i,j} A_{E,j}^{(p,0)}}{\left[J_{E,I}\right]_i}\right), \tag{S5}$$

where $\Gamma^{(p)} \stackrel{\text{def}}{=} \left\{i \in \{1, \cdots, 34\}\colon\ A_{E,i}^{(p,0)} = 1\right\}$. In other words, $A_{E,i}^{(p,0)}$ is not a solution to Eq. (S4) if $z > z^{(p)}$. It follows that the number of stationary attractors, $M^{(z)}$, decreases with $z$, because more and more solutions $A_{E,i}^{(p,0)}$ do not satisfy Eq. (S5) for large values of the scaling coefficient.

***S5 Model-based estimation of excitation-inhibition balance in cortical resting-state dynamics***

Given that our model included excitatory (E) and inhibitory (I) populations in each region, we finally examined which local E-I interaction patterns emerged from the best-fit model. We first investigated the balance between E and I currents received by either E or I populations in each region. The model predicts that during resting-state activity this balance is such that the total input current to each population (the signed sum of the E and I input currents) fluctuates relatively close to the threshold for firing (Fig. S5A). Consequently, the time-averaged spiking activity in both the E and I populations in each region (Fig. S5B) was at an intermediate level of 0.4 to 0.7, measured in a normalized scale in which zero indicates total silence and 1 indicates continuous firing always saturated at the maximal level. The overall E-I balance is further highlighted by the finding of a linear relationship between the total input current to the I population and the total input current to the E population in each cortical region (Fig. S5C), meaning that the regions with higher total input current to the I population have also a proportionally higher input current to the E population, which maintains E-I interactions balanced.

While there was a proportionality of the total currents received by E and I populations in the same region, some regions had a higher ratio of E/I input currents. In particular, regions in the DMN, which had high activity both in empirical and model fMRI time series, received a higher proportion of E input currents to the E population (Fig. S5D). The ratio of E/I input currents received by the I populations remained relatively stable across regions (Fig. S5D). Note that during the fitting procedure we did not attempt to achieve neither the linear relationship between the total input currents to the E-I populations, nor the balance of the mean firing rates described above. These properties emerged in the dynamics of our best-fit network.

We next investigated the functional consequence of the locally balanced nature of the E-I interactions predicted by our model. We computed the sensitivity of the response of the whole cortex to the application of an external perturbation current to all the populations. Results (Fig. S5E) indicate that the sensitivity is maximal around the value of 0 (unperturbed resting state activity) for the input current. This is because in the resting configuration the total input currents on each cortical region fluctuate around the firing threshold (see Fig. S5A), where weak perturbations in the stimulus intensity elicit the largest variations in activity. In sum, these results suggest the presence of a balanced nature of E-I local interactions in resting-state activity, and a possible advantage of high sensitivity to perturbation (and thus high stimulus information coding capabilities) of the resting-state configuration.

To highlight the importance of the E-I interactions predicted by the parameters of the best-fit model, we multiplied the weights of the local E to I and I to E anatomical connections by a global scaling coefficient $z$, so that the E-I populations are anatomically disconnected when $z = 0$, while they are connected by the best-fit weights when $z = 1$. We then studied the network dynamics as function of the scaling coefficient $z$. We found that both the linear relationship between total input currents to E and I populations and the sensitivity of cortical activity to the application of an external input current peaked at the $z \sim 1$ value, corresponding to the parameter values estimated by the model best fit to the empirical fMRI data. This suggests that the mouse brain at rest not only keeps E and I interactions balanced, but also that this balance is optimal for encoding of external stimuli.

| | Our Model | Undirected Model | Linear Model |
|---|---|---|---|
| $V^{\mathrm{thr}}$ | 1.4 | 1 | - 0.5 |
| $\sigma$ | 1.56 | 1.2 | 1 |
| $G_{E,E}$ | 0.17 | 0.18 | 0.01 |
| $J_{I,I}$ | - 0.5*[1,1…1] | - 0.5*[1,1…1] | - 0.5*[1,1…1] |
| $J_{I,E}$ | 5*[1,1…1] | 5*[1,1…1] | 5*[1,1…1] |
| $J_{E,I}$ | -0.3*[10, 4.5, 3, 8, 12, 5, 3, 1, 1, 1, 1, 1, 1, 1, 5, 2, 3] | -0.2*[5, 15, 17, 6, 10, 8, 8, 3, 1, 4, 1, 8, 6, 1, 6, 6, 5] | -0.0035*[8, 5, 3, 6, 11, 8, 3, 2, 1, 1, 1, 2, 1, 1, 4, 1, 3] |

**Table S1:** *Best-fit parameters*. In this table we list the best-fit parameters of the three whole-cortex models investigated in section 2.6: our model with directed anatomical connections and non-linear (threshold-based) activation function, the non-linear model with undirected connections between the excitatory populations, and finally the model with linear activation function and directed connections. For $J_{E,I}$, $J_{I,E}$ and $J_{I,I}$, each entry denotes the strength of the connections for a given region, numbered from 1 to 17 as in Fig 1A. For any given region, the parameters are the same across both hemispheres.

| Attractor Index | Decimal Representation | Attractor Index | Decimal Representation |
|---|---|---|---|
| 1 | 0 | 17 | 1188959372665946640 |
| 2 | 274877906960 | 18 | 1188968306197922840 |
| 3 | 9070970929680 | 19 | 1188977102290945560 |
| 4 | 18004502905880 | 20 | 2359886204879503360 |
| 5 | 26800595928600 | 21 | 2359886479757410320 |
| 6 | 36028797021061120 | 22 | 2359895275850433040 |
| 7 | 36029071898968080 | 23 | 2359904209382409240 |
| 8 | 36037867991990800 | 24 | 2359913005475431960 |
| 9 | 36046801523967000 | 25 | 3512807709553459200 |
| 10 | 36055597616989720 | 26 | 3512807984431366160 |
| 11 | 1152930300766978560 | 27 | 3512816505646481920 |
| 12 | 1152930575644885520 | 28 | 3512816780524388880 |
| 13 | 1152948305269884440 | 29 | 3512825714056365080 |
| 14 | 1188950301695016960 | 30 | 3512834510149387800 |
| 15 | 1188950576572923920 | 31 | 3801067085341132440 |
| 16 | 1188959097788039680 | | |

**Table S2:** *List of stationary attractors of our model.* In this table, we list the 31 stationary attractors of our model with the best-fit parameters. The attractors are sorted by ascending decimal representation. To recover the stationary activity patterns, these numbers have to be converted into the corresponding binary representations of length 68. For example, attractor 29 is 3512825714056365080 in base 10, which corresponds to the string $0011 \cdots 1000$ in base 2. In turn, this string represents the stationary activity pattern $\boldsymbol{A} = (0,0,1,1,\cdots,1,0,0,0)$. Attractors 1, 7, 11, 17, 23, 30, 31 are homotopic (i.e. $A_i = A_{i+17}$, $\forall i \in \{1,\cdots,17\}$ and $\forall i \in \{35,\cdots,51\}$), while the others exhibit distinct firing motifs in the two hemispheres. In this table we did not report the oscillatory attractors, since our algorithm detected a very large set of 34,877 oscillations (all having period $\mathcal{T} = 4$) over 100,000 repetitions of the network.

| **Attractor Index** | **Decimal Representation** |
|---|---|
| 1 | 0 |
| 2 | 9070970929680 |
| 3 | 1152930300766978560 |
| 4 | 1152930575644885520 |
| 5 | 1188959097788039680 |
| 6 | 3512807709553459200 |

**Table S3:** *List of stationary attractors of the undirected model.* As in Table S2, but for the model with undirected anatomical connections between the excitatory populations.

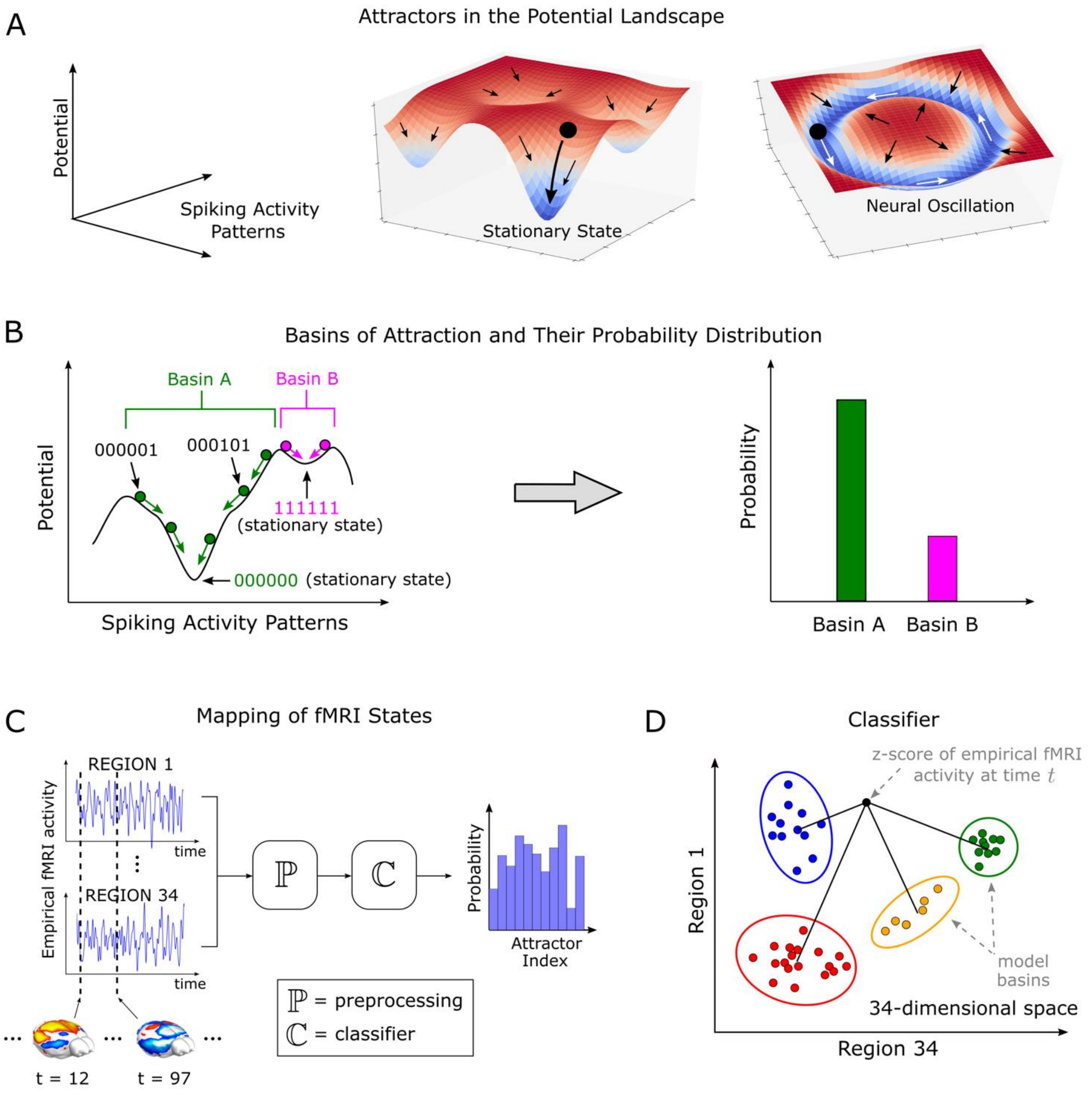


**Figure S1:** *Attractors and mapping algorithm*.

**A)** Intuitive interpretation of basins of attraction as valleys, and of attractors as minima, in an energy potential landscape. Note that our network model exhibits only two kinds of attractors, namely stationary states and neural oscillations.

**B)** An example of basins of attraction and their probability distribution in a network composed of 6 populations. As suggested by intuition, the probability with which a basin is visited in the long-time limit is directly proportional to its width and depth, and inversely proportional to its potential.

**C-D)** Mapping algorithm. The clusters in panel D represent the projections of the basins of attraction of the model-generated activity patterns into the space of the fMRI signals. At every time instant, the classifier labels the to-be-analyzed fMRI data point with the index of the closest cluster. Then, by counting the number of times each label has been used, the algorithm reconstructs the probability distribution of the basins of attraction. Finally, this distribution is compared with its model counterpart, which is obtained from the size of the clusters in panel D.

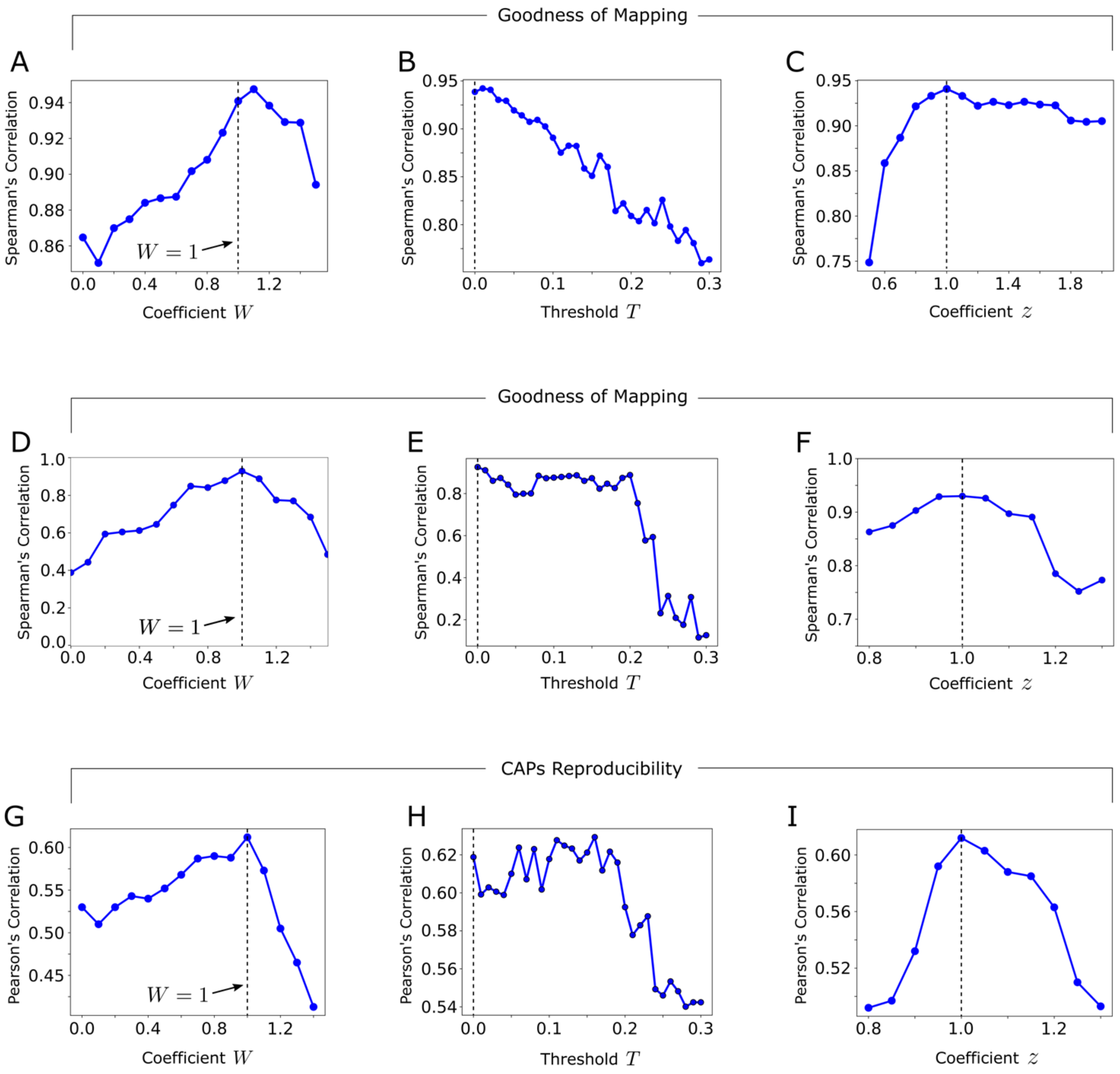


**Figure S2:** *Goodness of mapping and capability of the model to explain co-activation patterns.* **A)** Spearman's correlation between the distribution of the attractors reconstructed when the mapping algorithm embedded with the best-fit-model attractors is applied on the timeseries generated by the non-best-fit model, and the distribution of the embedded best-fit attractors, as a function of the global scaling coefficient $W$. **B)** As panel A, but as a function of the threshold $T$. **C)** As panels A, B, but as a function of $z$. **D)** Spearman's correlation between the distribution of the attractors reconstructed when the mapping algorithm embedded with the non-best-fit-model attractors is applied on the empirical timeseries, and the distribution of the embedded non-best-fit attractors, as a function of the global scaling coefficient $W$. **E)** As panel D, but as a function of $T$. **F)** As panels D-E, but as a function of $z$. **G-I)** Mean Pearson's correlation, calculated across the 6 CAPs of the mouse brain, between the empirical CAPs and those reconstructed through the model attractor dynamics. The model with the best-fit parameters ($W = z = 1$ and $T = 0$) exhibits the highest mean correlation.

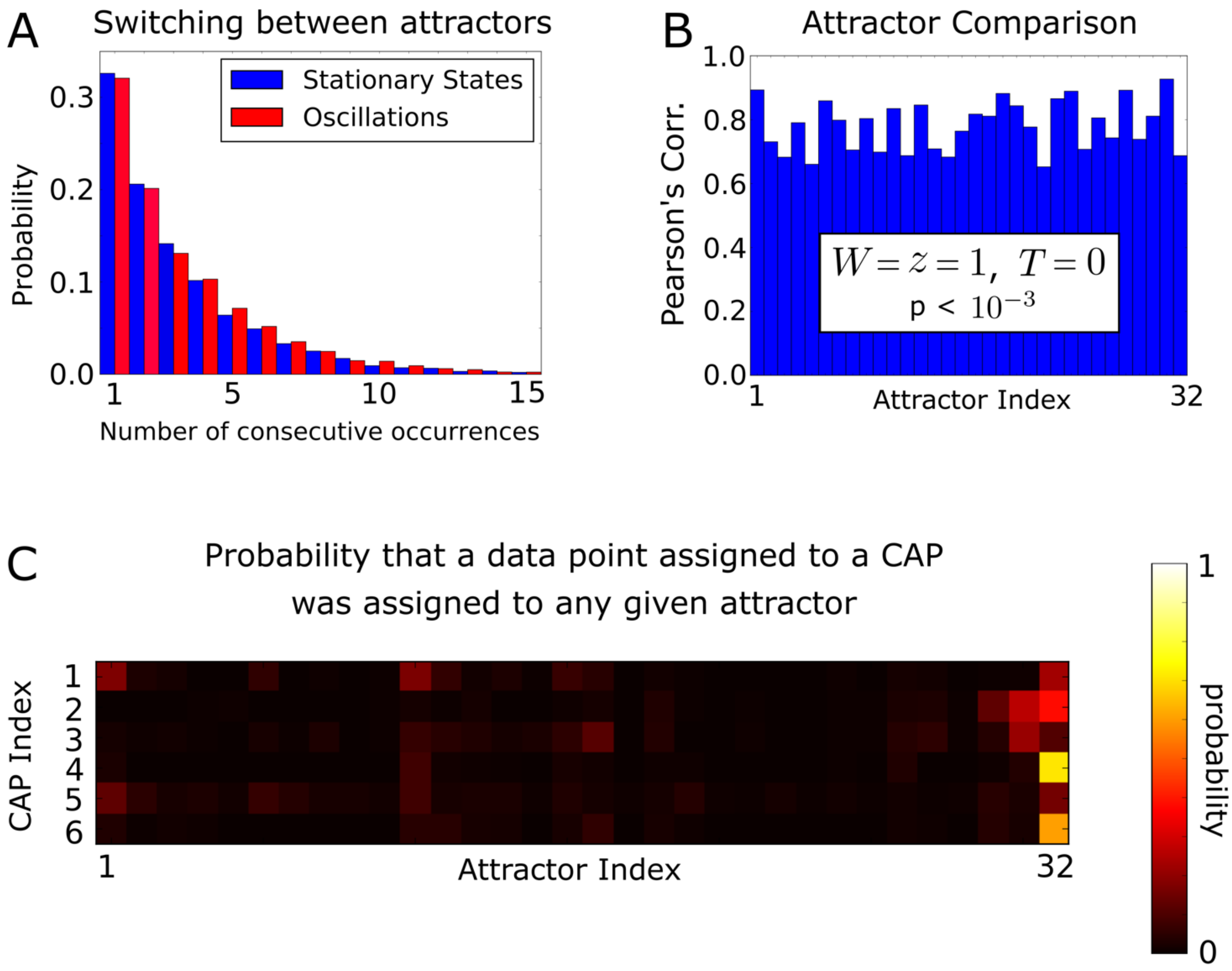


**Figure S3:** *Temporal and spatial properties of the model attractors dynamics, and reconstruction of CAPs from the spatial activation of each basin of attraction.*

**A)** Probability distribution of the number of consecutive time steps the activity pattern spends in any basin of the stationary attractors before visiting any basin of the oscillatory attractors (blue), and vice versa (red). **B)** Pearson's correlation for the model with the best-fit parameters ($W = z = 1$ and $T = 0$) between the mean z-score vectors of the model and empirical fMRI signals in each basin of attraction reconstructed by the mapping algorithm. **C)** Weights of the linear combinations of the model mean z-score vectors from which we reconstructed the topography of the 6 CAPs of the mouse brain for $W = z = 1$ and $T = 0$.

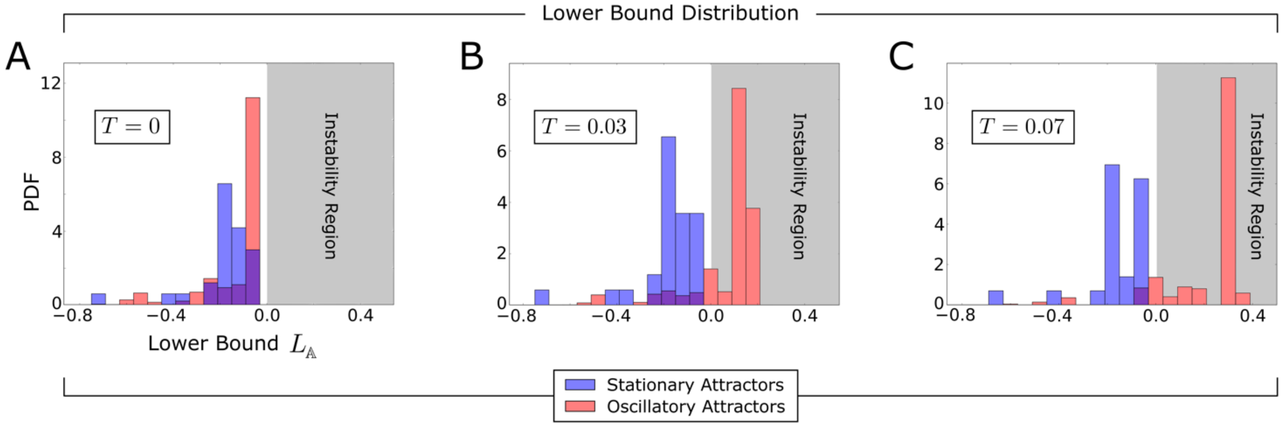


**Figure S4:** *Role of the strong vs weak anatomical connections*.
The figure reports, for different values of the anatomical threshold $T$, the distribution of values of $L_{\mathbb{A}}$ (see Eq. (S2)) of the stationary and the oscillatory attractors of our network model (blue and red bars, respectively).
**A)** Distribution for the unperturbed network ($T = 0$). All the attractors are stable.
**B-C)** The distribution of $L_{\mathbb{A}}$ is affected by the removal of the anatomical connections that are weaker than, e.g., $T = 0.03$ and $T = 0.07$. Note that $L_{\mathbb{A}}$, for most of the oscillatory attractors, becomes positive (shaded grey area), therefore those oscillations become unstable when the mouse is at rest (compare with the red curve in Fig. 6B). On the other hand, these panels show that the stability of the stationary attractors is not affected by the removal of the weakest anatomical connections (compare with the blue curve in Fig. 6B for $0 \leq T \leq 0.07$).

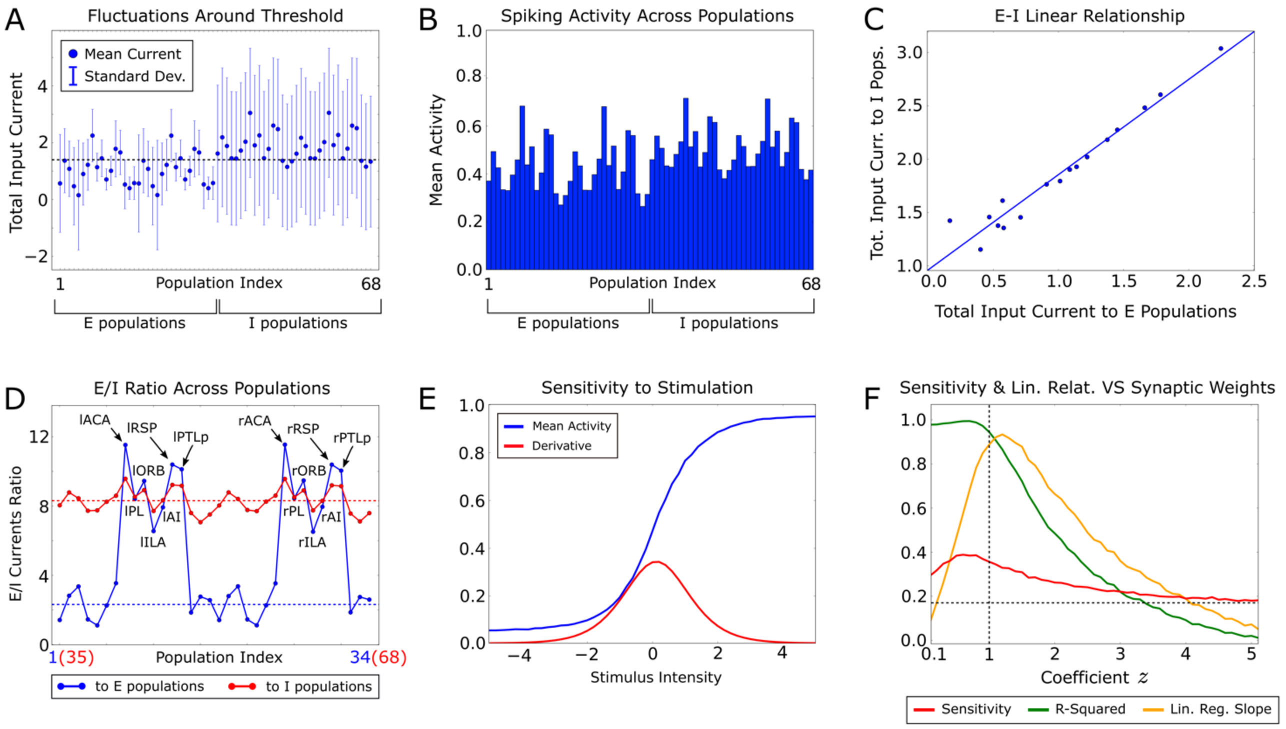


**Figure S5:** *Excitation-inhibition balance.*

**A)** Fluctuations of the total (i.e. E plus I) input currents to each cortical population. Excitation and inhibition balance each other, meaning that the E-I currents sum up to produce total currents fluctuating around the firing threshold $V^{\text{thr}}$ (dashed horizontal line).

**B)** Mean spiking activity of the cortical populations.

**C)** Linear relationship between the total currents to the E-I populations.

**D)** Ratio of the E-I components of the currents to each cortical population. The ratio is nearly constant for the inhibitory populations (red curve), while the excitatory populations of the DMN show a much larger ratio than the other excitatory subnetworks (blue curve).

**E)** Sensitivity of our model to external stimulation is maximal at rest, thereby suggesting a functional benefit of the balanced state. To evaluate sensitivity, we first computed the response to an external perturbation current to all the populations, averaged over the 10 simulation time steps following the stimulus, as a function of the perturbation strength, and then we quantified sensitivity as the derivative of the response with respect to the strength of the applied input current.

**F)** E-I linear relationship and its functional benefit, as a function of a global scaling coefficient $z$, which multiplies the synaptic weights between the E-I populations. Note that the sensitivity to stimulation is almost maximal for our model (i.e. for $z = 1$), and that the linear relationship between the total currents vanishes for large $z$.
